# One-Carbon Metabolic Reprogramming Stratifies Prognosis and Defines Distinct Biological States in IDH1-Mutant Gliomas

**DOI:** 10.64898/2026.09.07.749811

**Authors:** Mayank Bajaj, Roy Karnati

## Abstract

**Background:** IDH1 mutations in diffuse gliomas undergo epigenetic transformation and metabolic stress, through accumulation of 2-hydroxyglutarate. While these tumours generally exhibit a less aggressive phenotype, significant prognostic heterogeneity persists. The extent to which the One Carbon Metabolism (OCM) network is rewired in IDH1-MT gliomas and its contribution to this heterogeneity remain poorly understood.

**Methods:** We analysed the expression of 64 OCM genes using TCGA and CGGA transcriptomic datasets (n = 709). Composite OCM score, Cohen’s d, PCA and driver gene analysis were used to quantify effect sizes between IDH1-WT and MT cohorts. Prognostic relevance was assessed using Kaplan-Meier and Cox regression analyses. OCM metabolic rewiring and dependency were validated using *in vitro* U87 IDH1-WT and IDH1-R132H mutant glioblastoma cell line models. Furthermore, immune microenvironment interactions were evaluated using CIBERSORTx to dissect OCM-associated correlations.

**Results:** Using transcriptional profiling and targeted *in-vitro* validation, we reveal a distinct two-state metabolic phenotype governed by the OCM network. We demonstrate that while IDH1-MT tumors primarily rely on a suppressed metabolic phenotype, they are supported by compensatory mitochondrial pathways and an aggressive subset aberrantly hyperactivates specific proliferative OCM genes. This high aggregate OCM score was significantly associated with poor overall survival in the IDH1-MT cohort (CGGA HR = 1.61, p < 0.001), identifying an aggressive subset that predominates the favourable IDH mutation status. Functional validation showed aggressive IDH1-MT cells lose their baseline metabolic flexibility with changes in critical OCM genes and exhibit a strict auxotrophic dependence. Further, we also define that OCM rewiring was also associated with immune cell abundance, suggesting high OCM activity correlating with monocyte infiltration and low M2 macrophage polarisation.

**Conclusion:** Our findings offer a framework to identify high-risk patients and propose translational strategies for IDH1-MT tumors that may be refractory to standard IDH inhibitors. Therefore, the study presents a conceptual model for metabolic precision therapy defining OCM metabolic subsets that distinguish indolent, mitochondrial-dependent tumours from aggressive, proliferation-dependent phenotypes in IDH1-MT gliomas.

## 1. Introduction

Cancer cells show a distinct metabolic adaptation called Warburg effect, characterized by a preference to aerobic glycolysis from oxidative phosphorylation. Unlike healthy cells, where glucose is metabolised through glycolysis followed by entering tricarboxylic acid cycle and electron transport chain to generate ATP in presence of oxygen, cancer cells maintain high glycolytic activity even in the presence of sufficient oxygen (1). This phenomenon reflects a metabolic overflow that occurs when nutrient uptake exceeds immediate energy demands (2,3). Such metabolic reprogramming is a hallmark of cancer and is critical for sustaining cellular proliferation, biosynthesis, and survival under stress (4). Aberrations in oncogenes and tumour suppressor genes are also known to disrupt this equilibrium between proliferation and differentiation that directly influences the cellular metabolism and energy homeostasis (5,6).

Isocitrate dehydrogenase (IDH) is among the key metabolic enzymes implicated in this process, which catalyses the oxidative decarboxylation of isocitrate to α-ketoglutarate (α-KG). Somatic mutations in IDH1 (R132) and IDH2 (R172 or R140) have been identified in multiple malignancies, particularly in acute myeloid leukaemia (AML) and gliomas, with IDH1 mutations being far more frequent in low-grade gliomas (LGGs) (6,7). Mutant IDH enzymes lose their normal catalytic function and instead catalyse the reduction of α-KG to D-2-hydroxyglutarate (D-2HG), consuming NADPH in the process. The excessive production and thereby accumulation of D-2HG disrupts the TCA cycle by depleting key intermediates and energy substrates. Furthermore, loss-of-function mutations in D-2-hydroxyglutarate dehydrogenase (D2HGDH), the enzyme responsible for converting D-2HG back to α-KG, result in 2-hydroxyglutaric aciduria, which contributes to widespread epigenetic dysregulation, further promoting oncogenesis and tumour progression (8). Deficiency or mutations in D2HGDH cause the accumulation of D-2HG, which has also been linked to several tumour types, including gliomas, AML, and cartilaginous neoplasms such as chondrosarcomas (9,10). Elevated levels of D-2HG sensitise tumour cells to alkylating agents (11) and PARP1 inhibitors (12) by impairing DNA repair pathways. D-2HG blocks histone demethylation and causes genomic instability by compromising homologous recombination repair thereby enhancing tumour vulnerability (13). Beyond these functional changes, IDH mutations exert broad alternations in cellular metabolism, and therapeutic responses that differ compared to IDH1-WT (IDH1-WT) (14). IDH1 mutant (IDH1-MT) glioma cells exhibit increased uptake of glutamine and glutamate, utilising glutaminolysis to replenish TCA cycle intermediates and maintain NADPH balance. These cells also show altered amino acid metabolism, characterised by elevated levels of glycine, serine, and threonine, along with a reduction in aspartate, and N-acetylated amino acids (15,16). In addition to pathways involving glucose, glutamine, and acetyl-CoA metabolism, both α-KG and D-2HG play regulatory roles in several other metabolic processes, such as redox homeostasis and biosynthesis of lipids that require NADPH as a cofactor (17).

One-carbon metabolism (OCM), a critical network linking the folate and methionine cycles **(Figure 1)** to nucleotide synthesis, methylation reactions, and redox regulation, is one metabolism that is dysregulated in cancers due to widespread DNA and histone hypermethylation. Enzymes involved in the mitochondrial folate cycle, such as methylenetetrahydrofolate dehydrogenase 2 (MTHFD2) and serine hydroxymethyl transferase (SHMT), are reported to be significantly upregulated in several cancers, including hepatocellular carcinoma, colorectal cancer, breast cancer, and glioblastoma (18,19). This OCM stress is inherently linked to the IDH1 pathology through two key axes: (a) the depletion of NADPH, a cofactor essential for maintaining the thermodynamic flux in the OCM pathway, and (b) the epigenetic demands imposed by G-CIMP, a target of α-KG dependent dioxygenases, which drastically alters the consumption of methyl groups derived from OCM. Consistent with metabolomics analysis studies have shown decreased levels of the OCM amino acids, glycine and serine in IDH1-MT glioma compared to IDH1-WT tissue. These amino acids are precursors for one-carbon (1C) unit donation, and their deficit suggests a functional impairment or metabolic bottleneck within the OCM pathway of IDH1-MT cells. This impairment can be attributed to reduced NADPH to NADP ratio, which undermines the thermodynamic flux necessary for cytosolic serine synthesis and overall 1C unit transfer (20–23). The metabolic and epigenetic alterations induced by D-2HG extend beyond the tumour cell itself, shaping an immunosuppressive tumour microenvironment. IDH-MT gliomas typically show an immunologically quiescent or cold phenotype, characterised by significantly reduced infiltration of immune cells, including cytotoxic CD8+ T lymphocytes, CD4+ helper T cells, macrophages, and microglia, compared to IDH-WT gliomas (24,25). Furthermore, IDH-MT gliomas employ epigenetic changes to evade immune surveillance, including downregulation of natural killer (NK) cell ligands. This reduced host immune response is considered a contributing factor to the overall favourable prognosis associated with the IDH mutation (26–29).

**Figure 1.**
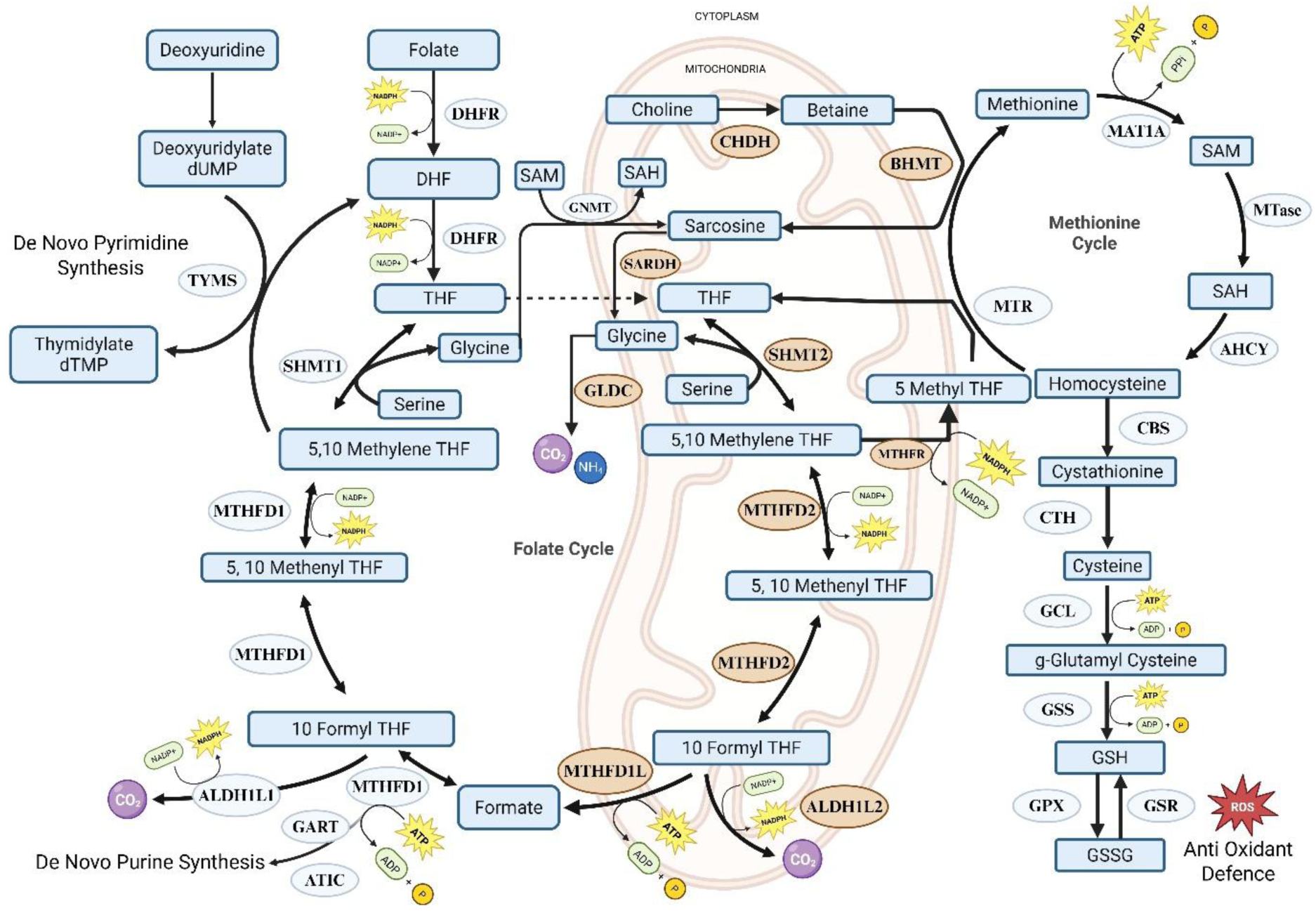
Overview of the compartmentalised one-carbon metabolic network and linked biosynthetic pathways. The network integrates nutrient inputs (folate, methionine, serine, and glycine) to support three fundamental cellular processes: (3) de novo synthesis of purines and pyrimidines (thymidylate) required for DNA replication; (2) generation of the universal methyl donor S-adenosylmethionine (SAM) for methylation reactions; and (3) production of glutathione (GSH) via the transsulfuration pathway for antioxidant defense against reactive oxygen species (ROS). The exchange of one-carbon units between the mitochondria and cytoplasm is mediated primarily by serine, glycine, and formate. Enzymes are depicted in ovals, and metabolites are shown in rectangular boxes. Cofactors (NADPH, ATP) indicate energy-consuming or redox-dependent steps.

Although, the altered OCM has been associated with differences in patient survival outcomes across several cancer types, including head and neck, colorectal, pancreatic, breast, lung, and paediatric brain tumours (30–32), and despite the established role of IDH mutations in reshaping global metabolism and epigenetic regulation, the extent to which these alterations reprogram OCM, while dictating prognostic heterogeneity and contribute to glioma aggressiveness within IDH1 mutant cohort remains insufficiently understood. While previous studies indicate a severe defect in the OCM network, the mechanistic relationship between the IDH1 mutation and the differential regulation of the OCM across the heterogeneous landscape of gliomas has not been comprehensively established (33–35). Thus, identifying OCM gene signatures could provide urgently needed biomarkers to distinguish indolent from aggressive disease, guide precision treatment selection for high-risk patients, and help segregate survival outcomes within the IDH1-MT cohort, independent of other markers such as TP53, TERT, or ATRX mutation status. This study, therefore, addresses the gap by evaluating OCM rewiring in IDH1-MT gliomas, elucidating its functional consequences, and linking it to differences in patient survival.

This study demonstrates that OCM is differentially regulated in gliomas and strongly linked to IDH1 mutation status, using gene expression profiles in the OCM network and overall signature-level activation across glioma grades and defined IDH1 subtypes, thereby reflecting IDH1 mutation biology. Further, the study investigates the biological outcomes defining the functional consequences of signature regulation specifically within IDH1-MT gliomas by characterising the immune environment associated with the signature, correlating individual OCM genes with immune cell infiltration, and correlating the global signature score with the IDH1-MT immune environment. The study further establishes two distinct biological states the in IDH1-MT gliomas regulated by OCM signature based on its association with patient survival outcomes. This study provides an integrated proof-of-concept framework demonstrating OCM as an essential vulnerability, opening new paths for targeted metabolic intervention and improving patient survival in IDH1-MT gliomas.

## 2. Results

### 2.1 IDH1-MT Gliomas Exhibit a Suppressed Proliferative One-Carbon Signature with Compensatory Upregulation of Mitochondrial 1C Source Pathways

Heatmap analysis of the expression profiles for all 64 curated OCM genes revealed transcriptional remodelling segregated by IDH1 mutation status in both the TCGA **(Figure 2A)** and CGGA **(Additional File 1, Figure S1)** datasets. This difference reveals differential metabolic dependency imposed by mutant IDH1 protein. The majority of OCM genes responsible for rapid cell growth, nucleotide synthesis, and methylation machinery were significantly downregulated in IDH1-MT cancers relative to IDH1-WT gliomas. This pattern also aligned with the established, less aggressive, slower-proliferating nature of IDH1-MT gliomas (41).

**Figure 2.**
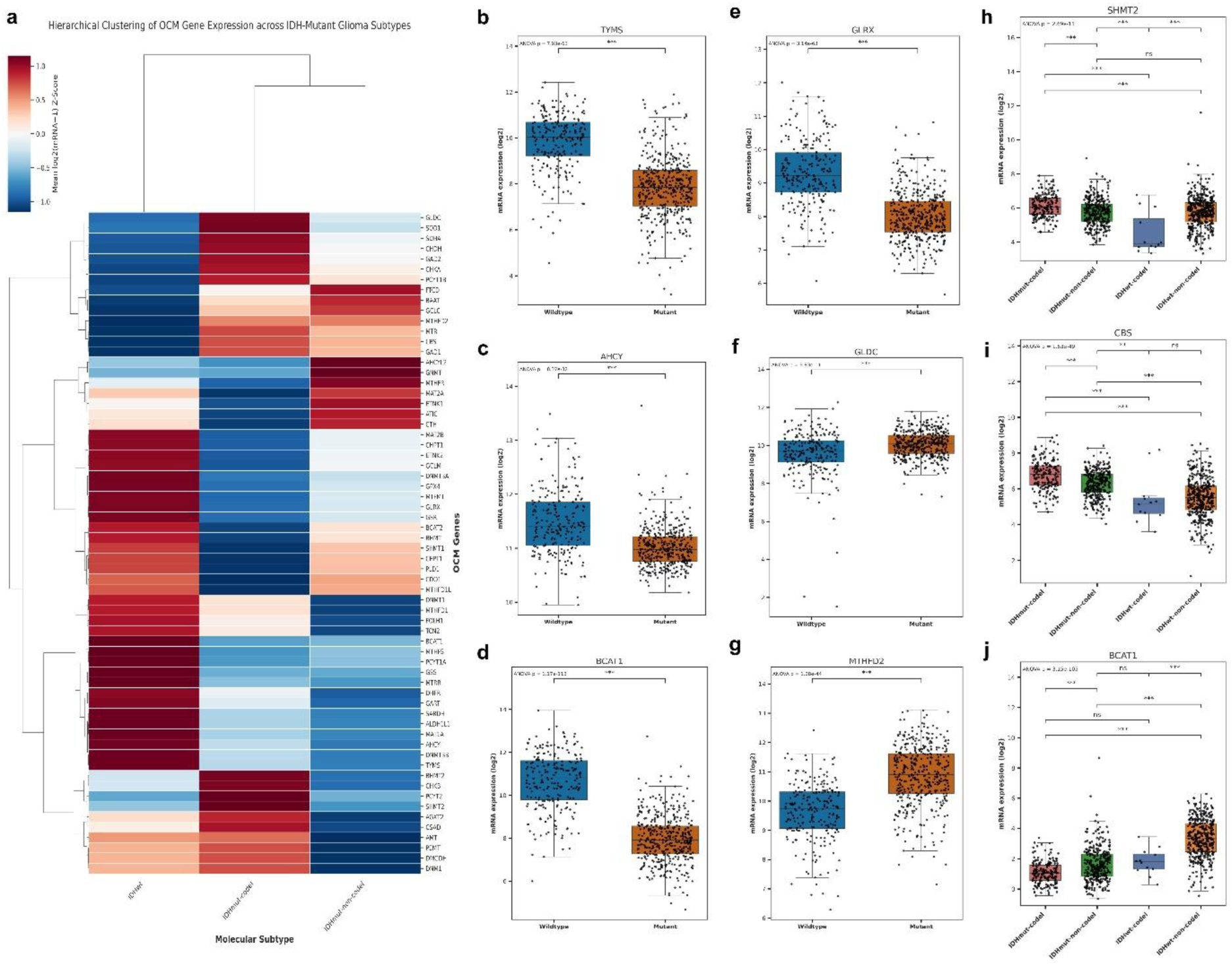
Differential Expression of One-Carbon Metabolism Reveals Divergent Proliferative and Compensatory Signatures Stratified by IDH1 Mutation and 1p/19q Status. **(a)** Hierarchical clustering heatmap of mRNA expression (Mean log2 Z-score) for OCM-related genes across molecular subtypes: IDH-wildtype (IDHwt), IDH-mutant with 1p/19q co-deletion (IDHmut-codel), and IDH-mutant without co-deletion (IDHmut-non-codel). Boxplots comparing the mRNA expression of key OCM genes between Wildtype and Mutant tumours, highlighting the expression of **(b)** TYMS, **(c)** AHCY, **(d)** BCAT1, **(e)** GLRX, **(f)** GLDC, and **(g)** MTHFD2. Gene expression profiles of **(h)** SHMT2, **(i)** CBS, and **(j)** BCAT1 were categorised across the distinct molecular subtypes to highlight the modulatory influence of 1p/19q co-deletion. Statistical significance was determined by ANOVA (*** P < 0.001; ns, not significant).

The suppression of the folate and nucleotide synthesis axis suggests reduced macromolecular demands characteristic of a lower mitotic rate. Key enzymes directly involved in nucleotide synthesis, such as TYMS (thymidylate synthase, required for *de novo* dTMP synthesis) (p=7.93e-63) and DHFR (dihydrofolate reductase) (p=2.02e-08), displayed high expression in IDH1-WT tumours but were markedly suppressed in IDH1-MT tumours. Similarly, GART (p=6.09e-05) and the cytosolic one-carbon carrier MTHFS (p=1.68e-60) showed significant downregulation in the MT cohort, reflecting diminished flux through the purine and folate cycles, respectively. The mitochondrial one-carbon intermediate recycler MTHFD1L that fuels the cytoplasmic folate cycle was also significantly downregulated in IDH1-MT tumours (p=5.34e-05). Enzymes responsible for generating or recycling the universal methyl donor, S-adenosylmethionine (SAM), and for maintaining DNA methylation were widely suppressed in IDH1-MT tumours, consistent with the global epigenetic effects of D-2HG. Specifically, AHCY (S-adenosylhomocysteine hydrolase, which regulates SAM/SAH ratio) (p=8.37e-32) and its recycling partner MTRR (p=1.11e-11) were significantly lower in IDH1-MT. The primary enzyme for SAM synthesis, MAT1A (p=1.25e-08), was also lower in MT tumours, suggesting a reduced overall capacity for SAM synthesis. Crucially, the *de novo* DNA methyltransferases DNMT3A (p=6.67e-47) and DNMT3B (p=2.98e-09) were significantly downregulated in IDH1-MT tumours, consistent with the generally lower proliferative rate of IDH1-MT tumours. However, the subtype remains characterised by the D-2HG-driven G-CIMP hypermethylated phenotype. **(Figure 2B-C, Additional File 1, Figure S2)**.

The aggressive IDH1-WT phenotype also relies heavily on catabolic pathways and robust redox capacity. Accordingly, catabolic enzymes such as BCAT1 (p=1.17e-113) and BCAT2 (p=6.16e-28) were drastically suppressed in IDH1-MT tumours, indicating severely restricted branched-chain amino acid catabolism, a hallmark of reduced metabolic plasticity observed in less aggressive cancer models (42). Furthermore, essential redox-regulatory genes, including GLRX (p=3.14e-63), GSS (p=1.1e-53), and GSR (p=4.3e-17), were significantly downregulated in IDH1-MT tumours, suggesting reduced oxidative stress or a fundamental metabolic shift away from glutathione-dependent detoxification pathways relative to IDH1-WT. Enzymes required for phosphocholine and lipid synthesis via the Kennedy pathway, such as CHPT1 (p=1.43e-24) and CEPT1 (p=1.7e-37), were also suppressed in IDH1-MT, suggesting reduced demand for membrane production consistent with slower proliferation **(Figure 2D-E, Additional File 1, Figure S2)**. The expression profile genes, such as ALDH1L1, CDO1, DNMT1, ETNK2, GPX4, MTFMT, PCYT1A, PLD1, SARDH, SHMT1, and TCN2, were in line with this lower expression trend seen in IDH1-MT tumours, thereby mutually defining a suppressed OCM signature.

The key enzymes for 1C generation exhibited substantial upregulated in the IDH1-MT tumours relative to the overall OCM transcriptional repression implying efforts towards preserving essential basal 1C flux for methylation or salvage pathways in the metabolically controlled IDH1-MT environment. The mitochondrial serine/glycine catabolism axis, which is crucial for 1C unit production, showed significant activation. MTHFR (folate methionine linkage component) (p=0.000257) and GLDC (glycine decarboxylase) (p=9.83e-11) were highly expressed in IDH1-MT tumours. This specific upregulation of mitochondrial 1C synthesis alongside high expression of the methyl donor detoxifier GNMT (p=0.0123), the mitochondrial folate pathway enzyme MTHFD2 (p=1.28e-44), and SHMT2 (p=0.00124), indicates that IDH1-MT cells rely on serine/glycine flux to generate one-carbon units. This could be a mechanism to sustain methionine cycling (via upregulated MTR) or to manage the resulting SAM/SAH imbalance caused by D-2HG accumulation **(Figure 2F-H, Additional File 1, Figure S3)**. Other genes showing high expression in the IDH1-MT cohort included CBS (p=2.44e-56), GCLC (1.09e-53), BHMT (p=0.000105), and CHDH (p=0.00235), which drive choline processing and are involved in the transsulfuration pathway to manage homocysteine levels or supply cysteine under stress. PCYT1B (p=4.83e-19) and BAAT (p=6.21e-18) were also highly expressed in IDH1-MT tumours, suggesting specific adaptations in lipid and amino acid conjugation pathways **(Figure S3).** Importantly, this divergent metabolic signature was largely corroborated in the independent CGGA dataset **(Additional File 1, Figure S4)**, with 20 genes exhibiting a similar trend, whereas 13 genes did not reach statistical significance.

Further, stratification of the datasets confirmed that OCM gene expression is also modulated by 1p/19q co-deletion. Genes demonstrating the strongest upregulation in IDHmut-codel tumours compared to IDH-WT subtypes included CBS (p=1.63e-49) and the 1C source genes AMT (p=0.00084), MTHFD2 (p=3.19e-37), GLDC (p=5.7e-15). The high expression of CBS in IDHmut-codel suggests that this specific subtype may have a heightened dependence on the synthesis of cysteine or other sulfur metabolites compared to IDHmut-non-codel and IDHwt groups. Conversely, highly proliferative markers such as BCAT1 (p=3.15e-05) and TYMS (p=1.17e-17) remained the lowest in IDH-MT tumours, irrespective of co-deletion status **(Figure 2I-J, Additional File 1, Figure S5)**.

### 2.2 IDH1 and Secondary Driver Mutations Synergistically Shift the OCM Transcriptome toward Compensatory 1C Metabolism

To evaluate the integrated effect of the IDH1 mutation on the overall metabolic landscape, a composite OCM signature score (mean Z-score) across all 64 genes was calculated for each sample. The overall OCM signature score was found to be much lower for IDH1-MT cohort (n=416) compared to the IDH1-WT cohort (n=293). This difference was highly significant (p = 4.228989e-39) and had a high biological effect size (Cohen’s d = −1.1520). Thus, the IDH1 mutation status maintained the strongest segregating factor even in the OCM **(Figure 3A)**. The PCA analysis on the 64 OCM genes revealed similar findings at the multivariate level, suggesting robust segregation of samples by IDH1 status. Majority of IDH1-WT samples clustered towards higher side of PC2 and intermediate sides of PC1, while being distinct from the IDH1-MT cluster. This confirmed that the combined expression pattern of OCM genes affects the overall segregation demonstrating that the OCM metabolic shift is a consequence of the IDH1 mutation **(Figure 3B)**.

**Figure 3.**
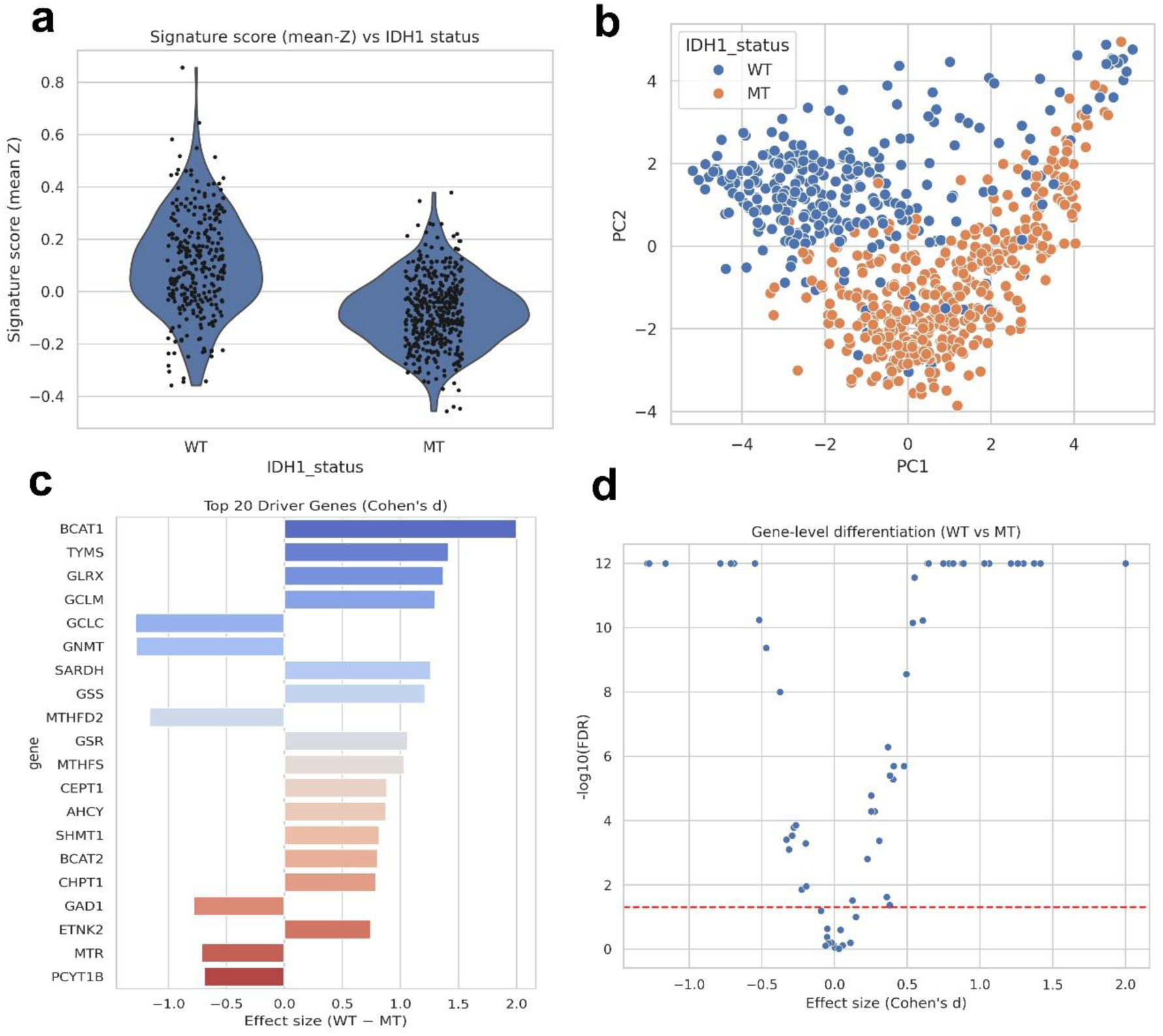
IDH1 mutation drives transcriptomic segregation and suppresses global one-carbon metabolism. **(a)** Violin plot comparing the distribution of the signature score (mean Z) between IDH1-WT and MT status (Cohen’s d = −1.1520, p = 4.228989e-39). **(b)** Principal Component Analysis (PCA) scatter plot illustrating the transcriptomic separation of tumour samples based on WT (blue) and MT (orange) IDH1 status. **(c)** Bar plot ranking the top 20 driver genes by effect size (Cohen’s d, WT - MT). Positive and negative values indicate genes downregulated and upregulated in IDH1-MT tumours, respectively. Colour blue, grey and red highlight sorting by magnitude in decreasing order. **(d)** Volcano plot displaying the statistical significance -log10(FDR) against gene-level differentiation effect size (Cohen’s d). The red dashed line denotes the threshold for statistical significance, highlighting the widespread transcriptional reprogramming driven by IDH1 status.

Additionally, to understand this global metabolic shift, analysis was performed to quantify the magnitude and direction of individual OCM gene contributions **(Table 1)** using Cohen’s d effect size. The genes showing strongest suppression in IDH1-MT (positive Cohen’s d) contribute the most to the higher signature score in IDH1-WT tumours. The effect sizes for key proliferative and catabolic enzymes were BCAT1 (d=2.000474), TYMS (d=1.416025), GLRX (d=1.370908), GCLM (d=1.297693), SARDH (d=1.259020), and GSS (d=1.211873). The marked differences in expression of these genes, particularly BCAT1, highlight the sharp distinction between the highly synthetic, energy-demanding IDH1-WT state and the metabolically suppressed IDH1-MT state. In contrast, the genes with the strongest upregulation in IDH1-MT (negative Cohen’s d) represent compensatory or essential maintenance pathways in the IDH1-MT. The most upregulated genes in MT were the 1C source and regulatory enzymes, including GCLC (d=−1.287133), GNMT (d=−1.275817), MTHFD2 (d=−1.162765), GLDC (d=−0.31277), and MTHFR (d=−0.29175). The bar plot ranking the top 20 drivers is represented in **Figure 3C**. This specific upregulation of 1C influx strongly supports the notion that the IDH1-MT tumours overcome the overall downregulation in proliferation-linked metabolism by improving their ability to feed the 1C cycle, favouring methylation over *de novo* synthesis to support rapid growth. The volcano plot, showing the distribution across all 64 genes, further confirmed that a large proportion of the OCM pathway is significantly regulated by IDH1 status **(Figure 3D)**.

**Table 1:** Statistical profiling and effect size analysis of the top 20 One Carbon Metabolism driver genes stratified by IDH1 mutation status. The table presents differential expression metrics comparing IDH1-WT and IDH1-MT tumours in the TCGA glioma cohort. Genes were selected based on their FDR-adjusted p-value to identify the most significant contributors to global metabolic reprogramming. Mean_WT & Mean_MT represent the average Z-score-normalised expression values for each gene in their respective cohorts. Delta: Indicates the absolute difference in mean expression (WT - MT). Cohen’s d: Quantifies the biological magnitude of the transcriptomic shift. A positive d denotes genes associated with the highly proliferative axis of IDH1-WT tumours (downregulated in IDH1-MT), while a negative d identifies key compensatory metabolic adaptations upregulated in IDH1-MT.

| Gene | Mean_WT | Mean_MT | Delta | Cohen_d | p value | FDR |
| --- | --- | --- | --- | --- | --- | --- |
| BCAT1 | 0.836055 | -0.58908 | 1.425139 | 2.000474 | 3.05E-74 | 1.95E-72 |
| TYMS | 0.681573 | -0.48024 | 1.16181 | 1.416025 | 7.86E-51 | 2.52E-49 |
| GCLM | 0.641686 | -0.45213 | 1.093819 | 1.297694 | 1.61E-50 | 3.44E-49 |
| GNMT | -0.63395 | 0.446685 | -1.08064 | -1.27582 | 3.78E-49 | 6.04E-48 |
| GLRX | 0.666745 | -0.46979 | 1.136534 | 1.370909 | 9.94E-49 | 1.27E-47 |
| GCLC | -0.63797 | 0.449513 | -1.08748 | -1.28713 | 1.99E-48 | 2.12E-47 |
| SARDH | 0.627942 | -0.44245 | 1.07039 | 1.25902 | 1.55E-44 | 1.42E-43 |
| GSS | 0.610703 | -0.4303 | 1.041005 | 1.211873 | 1.84E-44 | 1.47E-43 |
| MTHFD2 | -0.59218 | 0.417247 | -1.00942 | -1.16276 | 6.18E-42 | 4.40E-41 |
| GSR | 0.552818 | -0.38952 | 0.942334 | 1.063255 | 1.13E-33 | 7.21E-33 |
| MTHFS | 0.538585 | -0.37949 | 0.918072 | 1.028717 | 1.15E-32 | 6.66E-32 |
| CEPT1 | 0.476358 | -0.33564 | 0.812 | 0.885394 | 2.30E-27 | 1.22E-26 |
| BCAT2 | 0.438238 | -0.30878 | 0.74702 | 0.802868 | 8.47E-25 | 4.17E-24 |
| AHCY | 0.471827 | -0.33245 | 0.804276 | 0.875393 | 2.42E-24 | 1.11E-23 |
| BAAT | -0.31025 | 0.218599 | -0.52884 | -0.54738 | 1.04E-23 | 4.44E-23 |
| SHMT1 | 0.444226 | -0.313 | 0.757227 | 0.815598 | 2.91E-23 | 1.17E-22 |
| GAD1 | -0.42958 | 0.302683 | -0.73226 | -0.78461 | 2.15E-22 | 8.11E-22 |
| MTR | -0.39565 | 0.278776 | -0.67443 | -0.71457 | 2.13E-21 | 7.56E-21 |
| CHPT1 | 0.430878 | -0.3036 | 0.734474 | 0.787331 | 1.72E-20 | 5.80E-20 |
| ETNK2 | 0.411239 | -0.28976 | 0.700998 | 0.746451 | 2.26E-19 | 7.24E-19 |

Subsequently, to evaluate whether the IDH1-driven metabolic state is stable or responsive to secondary oncogenic alterations, we investigated the influence of TP53, ATRX, and TERT mutations across median-stratified score groups (high vs. low) and the continuous distribution of OCM signature scores by mutation status. TP53 exhibited the highest overall mutation burden, with a comparable distribution between the high (∼43%) and low (∼45%) score groups. Similarly, signature scores were alike between TP53-mutated and wildtype samples. However, a distinct variation in ATRX mutation status was observed, with ATRX alterations more prevalent in the low OCM score group (36%) than in the high score group (23%), and tumours with ATRX mutations showed a noticeable downward shift in their overall score distribution **(Additional File 1, Figure S6)**. TERT mutations were exceedingly rare across the entire cohort (<1%) and demonstrated no association between the two groups in either analysis. Together, these findings suggest that while TP53 mutations represent a common baseline event independent of the OCM score, a lower score is associated with an enrichment of ATRX genetic alterations.

The Venn diagram **(Figure 4A)** revealed substantial overlap among OCM genes driven by IDH1, TP53, and ATRX status. 30 OCM genes were significantly regulated by all three major drivers, highlighting that core OCM metabolism is governed by the central genetic alterations defining diffuse gliomas. Only 10 genes were uniquely sensitive to IDH1 mutation status **(Additional File 2, Table S2)**, while only 7 were co-regulated by IDH1 and ATRX, and 4 were co-regulated only by IDH1 and TP53. This vast convergence confirms that the OCM pathway is a highly integrated metabolic effector network for these oncogenic pathways. Further analysis of the effect size of TP53 and ATRX mutations showed that these drivers primarily alter the 1C source and regulatory genes that are already shifted by IDH1 status. TP53 mutation resulted in the highest upregulation of MTHFR (Cohen’s d ∼1.19), an enzyme central to folate-mediated 1C unit flow into the methionine cycle **(Figure 4B)**. Genes involved in 1C compensation, such as MTHFR, GCLC, GNMT, MTHFD2, BAAT and CTH, were also promoted by TP53 mutation. This suggests that TP53 loss or mutation synergistically enhances the 1C compensatory mechanism regulated by IDH1-MT. Conversely, SARDH, ALDH1L1, BCAT1, TYMS, MAT1A and GSS were among the genes suppressed by TP53 mutation, mirroring the general suppression of proliferative pathways observed in IDH1-MT tumours **(Additional File 2, Table S3)**. The ATRX mutation profile also largely resembled that of TP53, suggesting co-functional regulation of the OCM pathway **(Figure 4C)**. MTHFR (Cohen’s d ∼1.08) showed the greatest positive effect, followed by GNMT, GCLC, MTHFD2, SHMT1, BAAT, CTH, and AHCYL2. Similarly, SARDH, BCAT1, and GSS were the most suppressed genes in ATRX mutant tumours, followed by AHCY, TYMS, ALDH1L1, DHFR, MTHFS, and GART **(Additional File 2, Table S4)**. This dual regulation by TP53 and ATRX confirms that maintaining essential 1C supply and suppressing aggressive proliferation markers are shared metabolic requirements for these common IDH1-MT co-mutations. Overall, samples with ATRX or TP53 mutations tended to have a lower OCM signature score than their non-MT counterparts across the entire cohort. The TERT mutation fraction did not differ significantly between high- and low-OCM signature score groups.

**Figure 4.**
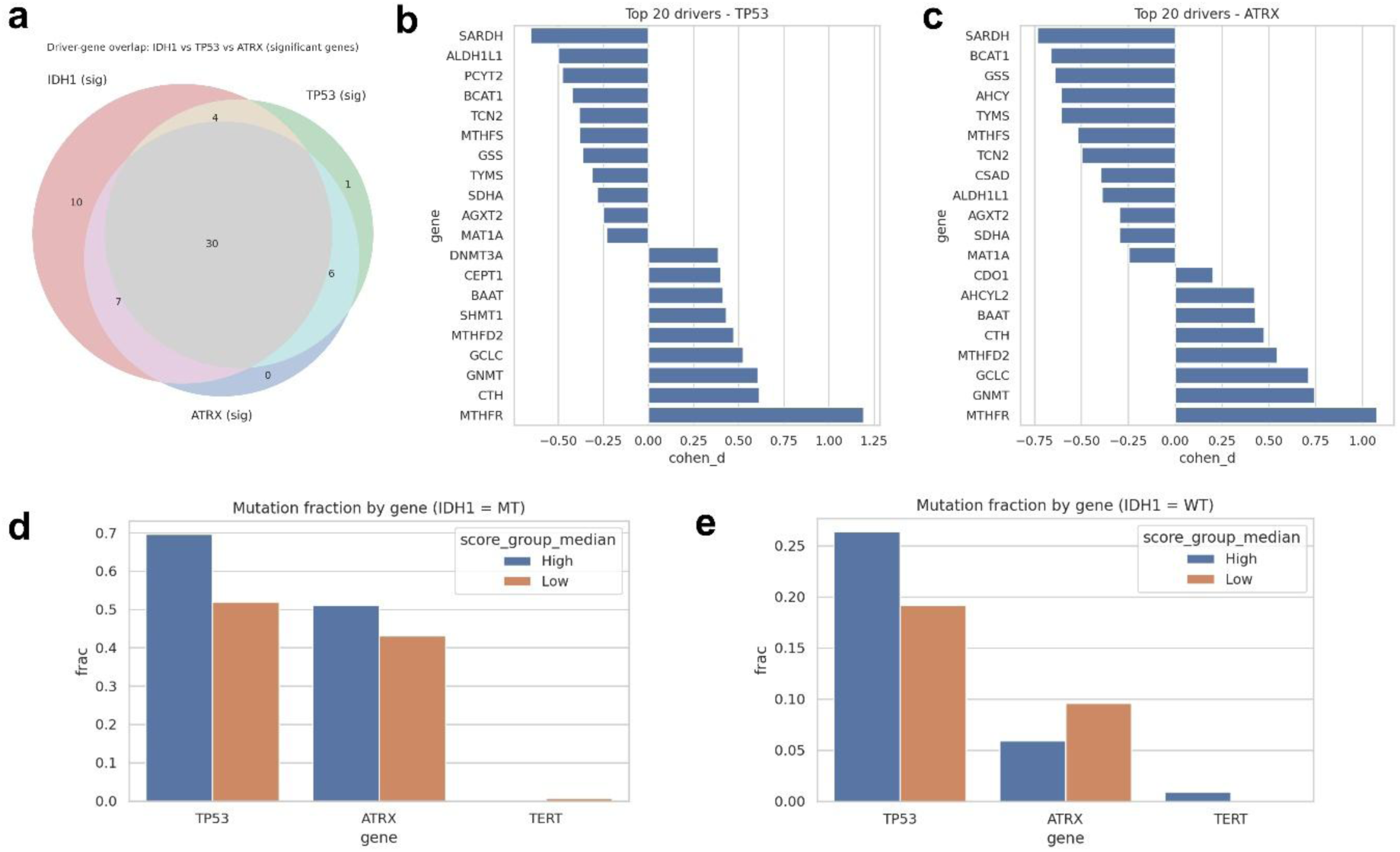
TP53 and ATRX co-mutations with IDH1-driven metabolic shifts define highly aggressive glioma sub-populations. **(a)** Venn diagram illustrating the driver-gene overlap among significant genes across IDH1, TP53, and ATRX, revealing a core set of 30 genes shared across all three genomic contexts. Bar plots ranking the top 20 OCM driver genes based on Cohen’s d effect sizes (Δ = MT − WT) associated with **(b)** TP53 and **(c)** ATRX. Bar plots displaying the mutation fraction by gene (TP53, ATRX, TERT) stratified by High (blue) and Low (orange) score group medians in the **(d)** IDH1-MT cohort and **(e)** IDH1-WT cohort.

We further stratified the influence of TP53 and ATRX by IDH1 status. Within IDH1-MT tumours, a high OCM signature score (above the median) was associated with a higher fraction of both TP53 mutations (∼70% in high score vs ∼ 52% in low score) and ATRX mutations (∼51% in high score vs ∼43% in low score). However, within IDH1-WT tumours, a high OCM signature score correlated with a higher fraction of TP53 mutations (∼ 26% vs ∼19% in low score) while a lower fraction of ATRX mutations (∼6% vs ∼9% in low score) **(Figure 4D-E)**. This divergent pattern indicates that a TP53 mutation enhances metabolic activity in aggressive IDH1-WT, whereas an ATRX mutation appears to restrict the highly proliferative OCM transcriptome, which is otherwise characteristic of IDH1-WT tumours. These results indicate that while the IDH1 mutation sets the global ceiling for OCM activity, secondary driver mutations, particularly TP53 and ATRX, fine-tune the metabolic profile, thereby promoting compensatory 1C-source pathways that define unique metabolic dependencies across glioma molecular subtypes. This finding of a convergence between high OCM signature score and the most aggressive genetic profiles (high TP53 and ATRX fraction) in IDH1-MT gliomas might provide a biological mechanism explaining the prognostic heterogeneity.

### 2.3. Transcriptional Reprogramming in U87 IDH1-MT Cells Confers High-Risk Metabolic Dependencies on Exogenous One-Carbon Units

To functionally validate the transcriptional rewiring observed in the tumour cohorts, the relative expression of core OCM genes was evaluated using RT-qPCR utilising U87 IDH1-WT and U87 IDH1 R132H-MT glioblastoma cell lines. The U87 parental cell line served as an inherently aggressive, primary Grade IV glioblastoma model. The targeted integration of the *IDH1* R132H mutation into this background (U87-MT) provided a high-risk subset model. Crucially, while IDH1 mutations are indicative of less aggressive phenotypes, the U87-MT line does not capture a lower-grade astrocytoma phenotype and functionally corresponds to secondary glioblastoma.

Consistent with the global transcriptional suppression of redox genes observed in the patient cohorts, the IDH1-MT cells exhibited significant downregulation of GLRX **(Figure 5A),** validating the fundamental shift away from specific glutathione-dependent detoxification pathways in the presence of the IDH1 mutation. Furthermore, the *in vitro* model successfully confirmed the compensatory upregulation of alternative 1C and transsulfuration pathways. Results show a marked upregulation of CBS, MTHFD2, MTR, and MTHFR **(Figure 5B-E),** confirming a conserved biological reliance on these axes to maintain 1C flux under IDH1 mutant conditions. Although IDH1-MT patient cohorts showed a general suppression of proliferative and methylation markers, the U87-MT cells show significantly elevated expression of TYMS, SHMT1, DNMT3B, ALDH1L1, DHFR, and MAT1A relative to wildtype cells **(Figure 5F-K)**. This transcriptional profile may mirror the specific high-risk subset identified earlier. This suggests that within the aggressive genetic subset, IDH1-MT cells override proliferative OCM pathways and DNA methyltransferases, thereby sustaining rapid growth and overcoming the indolent IDH1 phenotype. The lack of GLDC **(Figure 5L)** upregulation in IDH1-MT cells further suggests that specific components of the compensatory signature may depend on nutrient deprivation, which was investigated further.

**Figure 5.**
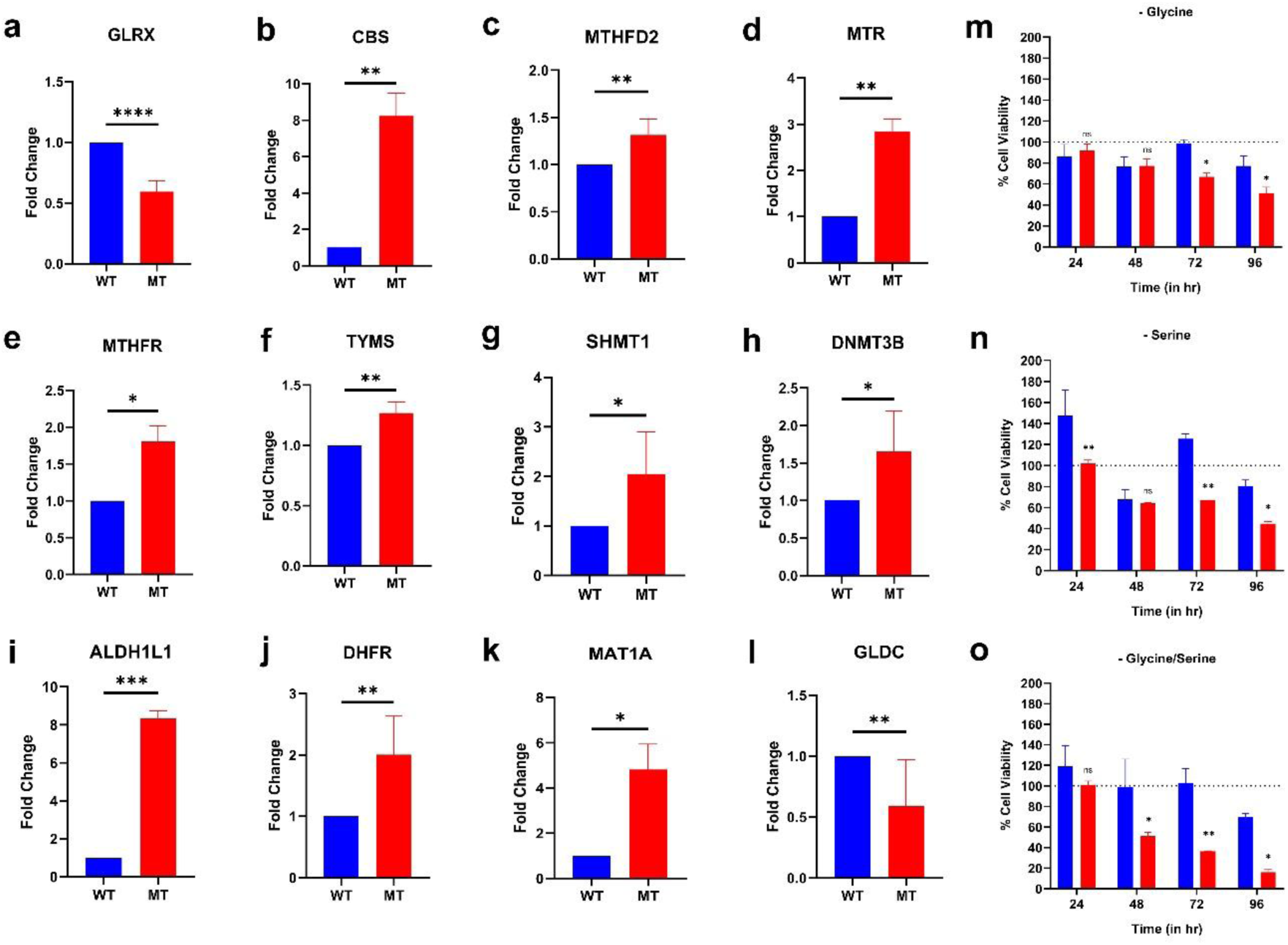
IDH1-mutant cells exhibit high-risk metabolic reprogramming and undergo apoptosis upon synergistic deprivation of serine and glycine. Bar charts showing the fold change expression and functional validation in IDH1-WT and MT cells for **(a)** GLRX, **(b)** CBS, **(c)** ALDH1L1, **(d)** MTHFD2, **(e)** MTR, **(f)** MTHFR, **(g)** TYMS, **(h)** SHMT1, **(i)** DNMT3B, **(j)** DHFR, **(k)** MAT1A, and **(l)** GLDC illustrating suppression of basal redox gene alongside the compensatory upregulation of transsulfuration, and one carbon axis. Elevated expression of proliferative markers suggests a high-risk secondary glioma aggressive IDH1-MT subset. RT-qPCR expression is normalised to housekeeping controls (β-actin). Error bars represent ± SEM of three independent replicates. Starvation assays monitoring percentage cell viability over 96 hours for WT and MT cells under (m) - Glycine, (n) - Serine, and (o) - Glycine/Serine deprivation conditions, demonstrating the heightened functional dependency of IDH1-MT cells on exogenous one-carbon units. All viability data represent mean ± SD from two independent biological replicates (* P < 0.05, ** P < 0.01, *** P < 0.001 Student’s t-test).

To evaluate the cellular reliance on exogenous 1C units, we next subjected U87-WT and U87-MT cells to deprivation of serine and glycine for 96 hours. Removing exogenous serine and glycine revealed a profound, targetable metabolic vulnerability in the IDH1-MT background. Cell viability over 96 hours revealed a divergence in metabolic plasticity between the two cell line models. Under only serine deprivation (-S), U87-WT cell viability surged at 24 hours, exhibiting an acute compensatory response, suggesting rapid upregulation of biosynthetic pathways while successfully adapting this restriction to maintain the viability by 72 hours. In sharp contrast, the U87-MT cells demonstrated a strict auxotrophic phenotype. The mutant cells failed to undergo metabolic rescue and exhibited a progressive, time-dependent collapse in cell viability across all starvation conditions. This functional defect was maximally seen under synergistic deprivation of both glycine and serine (-G/S). The IDH1-MT cells showed complete population collapse with a severe lethal endpoint of 21.58% viability **(Figure 5M-O)**. While WT cells temporarily survived this dual restriction until 72 hours by entering a protective state of metabolic quiescence, they too eventually declined by 96 hours. Thus, the functional assays confirm that, while the aggressive IDH1-MT subset aberrantly upregulates the proliferative 1C machinery, it completely loses baseline metabolic flexibility, becoming dependent on exogenous serine and glycine to maintain one-carbon flux and prevent cell death.

Taken together, the observed IDH1-driven transcriptional remodelling of the OCM signature, characterised by a fundamental suppression of proliferation-linked pathways alongside upregulation of compensatory axes, implicates the OCM signature as a determinant of clinical behaviour. The metabolic shift, representing the strongest molecular segregation (by IDH1 status), establishes a biological foundation for predicting prognosis. To further translate these profound metabolic distinctions into clinical findings, we next evaluated the prognostic utility of the derived OCM transcriptional signature across the heterogeneous landscape of gliomas.

### 2.4 The Global OCM Combined Score Independently Predicts Poor Survival Across Glioma Subtypes

To assess the translational utility of the differential OCM regulation, we computed a singular OCM combined score derived from the average Z-score of all 64 1C metabolic genes. This score was evaluated for its prognostic value across pan-glioma, IDH1-WT, and IDH1-MT cohorts in the CGGA and TCGA datasets.

In the pan-glioma cohort, the OCM combined score was confirmed as an extremely potent predictor of survival, validating the concept that high overall OCM pathway activation is characteristic of highly aggressive glioma regardless of IDH status. In the CGGA cohort, a high OCM combined score was significantly associated with poorer overall survival (HR = 1.79, 95% CI [1.51-2.11], Cox p = 7.49e-12) **(Figure 6A)**. This dramatic prognostic segregation was even stronger in the TCGA pan-glioma cohort (HR = 5.10, 95% CI [3.76-6.92], Cox p = 1.08e-25) **(Additional File 1, Figure S7)**, highlighting that the integrated OCM metabolic state accurately reflects tumour aggressiveness in the overall glioma population. In the aggressive IDH1-WT cohort, high OCM activity was associated with a marginally reduced survival. In CGGA, the high score group showed an HR of 1.28 (95% CI [1.03-1.59], Cox p = 0.0248), and the TCGA cohort showed an HR of 1.37 (95% CI [1.00-1.88], Cox p = 0.0511) **(Figure 6B)**. While less potent than in the pan-glioma setting, this confirms that OCM activation contributes to poor prognosis even in the aggressive IDH1-WT background.

**Figure 6.**
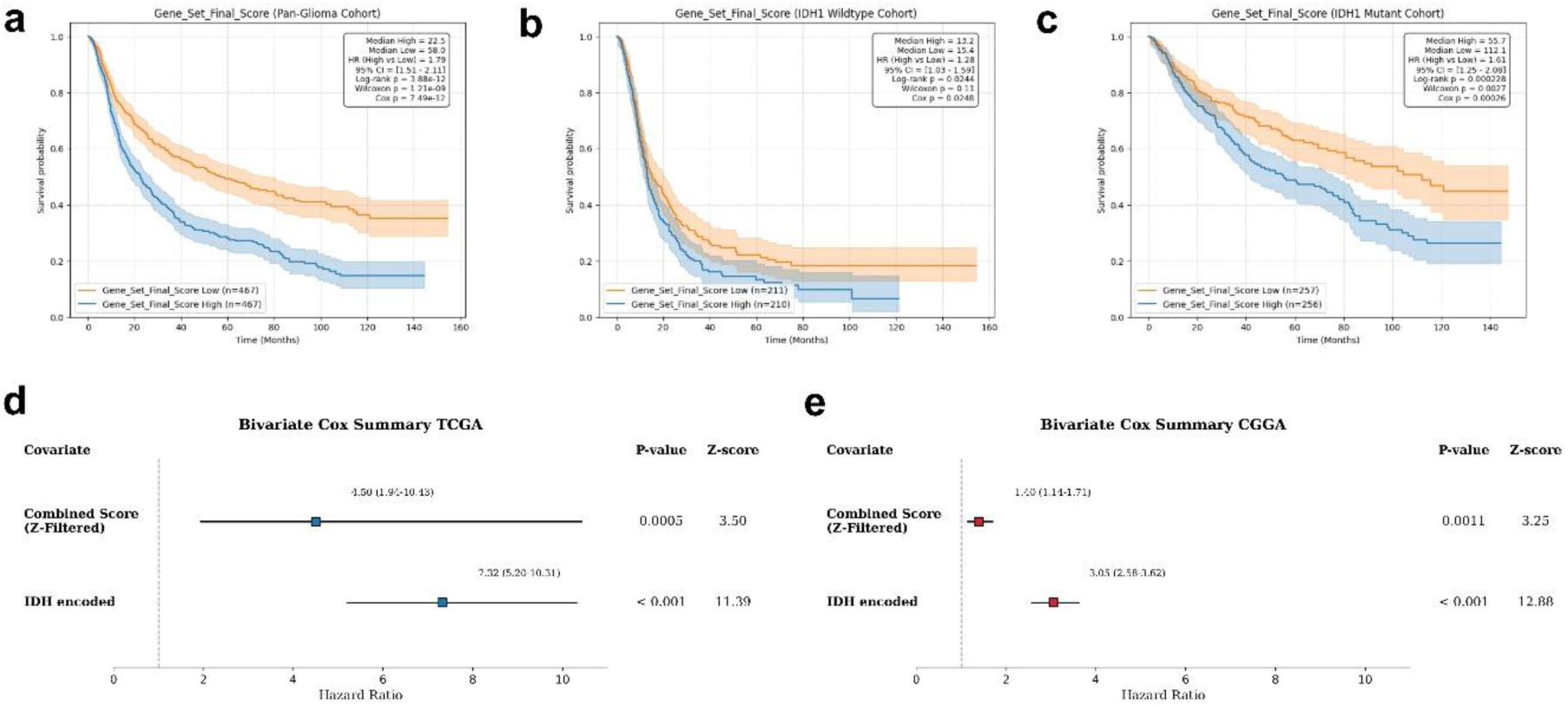
The OCM combined score is an IDH status independent robust prognostic marker in gliomas. Kaplan-Meier survival probability estimates stratifying patients (CGGA) into high-risk (orange) and low-risk (blue) groups based on the combined score across **(a)** the Pan-Glioma Cohort, **(b)** the IDH1 Wildtype Cohort, and **(c)** the IDH1 Mutant Cohort. Shaded regions represent 95% confidence intervals. Forest plots summarising bivariate Cox proportional hazards regression analyses using combined score and IDH encoded covariates in both **(d)** TCGA and **(e)** CGGA cohorts. The score retains significant prognostic value independent of IDH mutation status. Squares represent the hazard ratio, and horizontal lines indicate the 95% confidence interval.

The OCM signature also successfully partitioned the generally indolent IDH1-MT cohort into distinct survival subgroups. In the CGGA cohort (n=513), tumours exhibiting a high OCM combined score demonstrated significantly worse survival (HR = 1.61, 95% CI [1.25-2.08], Cox p = 0.00026). This finding demonstrates that OCM activity is a major determinant of prognostic heterogeneity in IDH1-MT disease and identifies an aggressive metabolic subset whose survival trajectory is abrogated despite the favourable IDH1 mutation **(Figure 6C)**. This finding suggests that high OCM transcriptional flux overrides the survival benefit typically conferred by the IDH1 mutation. However, this effect showed cohort-specific divergence where prognostic significance was attenuated in the TCGA IDH1-MT cohort (HR 1.26 (95% CI [0.80-1.98], Cox p = 0.32). This divergence can be attributed to factors such as reduced functional effects and reduced statistical power of 13 detrimental and 4 protective key prognostic genes in the TCGA cohort. For example, key proliferative markers that were highly detrimental in CGGA (e.g., TYMS HR 2.42, p = 3.28e-11; MTHFR HR 2.46, p = 2.22e-11) exhibited substantially reduced or non-significant prognostic power in the TCGA IDH1-MT cohort (TYMS HR 1.24, p = 0.359; MTHFR HR 1.24, p = 0.358), thus weakening the overall combined score’s ability to segregate survival clearly. This overall lack of statistical power for individual aggressive OCM genes in the TCGA IDH1-MT cohort suggests that the metabolic shift necessary to drive aggressive behaviour in this cohort was less marked and less closely linked to survival outcomes than in the CGGA cohort. The prognostic ability of the combined score in the IDH1-WT subset also shows a similar trend in the TCGA dataset. This general drop in prognostic precision for the aggregate score in TCGA is likely driven by the distinct clinical and demographic variations, ancestry-specific genetic architectures, follow-up duration, or treatment protocols inherent in the TCGA data. While the CGGA explicitly captures an East Asian patient cohort with a high proportion of recurrent tumours, the TCGA dataset is heavily skewed toward treatment-naive primary tumours from a predominantly white population (∼85%), contributing to the attenuated metabolic signals relative to those in the CGGA data.

To address this further, we employed a bivariate Cox regression model in pan glioma dataset and confirmed that the prognostic value of the OCM score is independent of the dominant IDH status prognostic factor. In both independent datasets, the OCM Combined Score maintained a significant prognostic association when controlled for IDH mutation status. Consistent with our results, the Combined Score remained statistically significant in the TCGA cohort, with a coefficient of 1.504 and an HR of 4.498 (p = 0.000459). Similarly, the Combined Score had a coefficient of 0.3337 and an HR of 1.396 (p = 0.001147) in the CGGA cohort **(Figure 6D-E)**. These results demonstrate that the OCM transcriptional signature captures biological information likely related to underlying proliferative potential or redox stress, and is statistically independent of the primary genetic stratification defined by the IDH mutation status.

### 2.5 Divergent OCM Pathway Activation Drives Prognostic Heterogeneity in IDH1-MT Gliomas

Given the significance of the combined score within the IDH1-MT CGGA cohort, we further investigated the prognostic role of individual OCM genes to identify metabolic components correlating with survival outcomes in IDH1-MT tumours. This analysis revealed two distinct metabolic profiles within the IDH1-MT subset: those activating proliferative pathways (associated with poor prognosis, HR > 1.0) and those robustly supporting compensatory pathways (associated with favourable prognosis, HR < 1.0).

#### 3.5.1 Hyperactivation of Proliferative OCM Genes Drives Aggressive Phenotypes in IDH1-MT Tumours

The globally suppressed OCM genes in IDH1-MT tumours, associated with proliferation and biosynthesis, emerged as poor prognostic markers when their expression was aberrantly high in the IDH1-MT cohort. This reflects a metabolic deviation towards an aggressive phenotype, essentially suggesting the prognostic advantage conferred by the *IDH1* mutation. Among the most powerful detrimental markers identified in the CGGA IDH1-MT cohort were TYMS (HR=2.42, p=3.28e-11), SHMT1 (HR=2.49, p=1.25e-11), MTHFR (HR=2.46, p=2.22e-11), GLRX (HR=2.14, p=1.08e-08), BAAT (HR=2.12, p=1.07e-08), BCAT1 (HR=1.93, p=6.21e-07), DNMT3B (HR=1.92, p=6.06e-07), AHCY (HR=1.63, p=0.000153), GART (HR=1.35, p=0.0192) **(Figure 7A-C, Additional File 1, Figure S8)**. The prognostic significance of SHMT1, TYMS, AHCY, BCAT1, and GART is important as they are markers of highly active nucleotide biosynthesis. Furthermore, MAT2B (HR=1.61, p=0.0444), MTHFS (HR=1.68, p=0.0274), and MTRR (HR=2.30, p=0.000541) also showed highly aggressive prognostic behaviour in TCGA IDH1-MT tumours, underscoring the critical role of the hyperactive methionine and folate cycles in promoting disease aggressiveness in this specific context. The upregulation of these proliferative genes, despite the generally suppressed global signature, indicates that a high proliferative drive mediated by OCM gene activation constitutes a major risk factor within IDH1-MT disease. The detrimental roles of SHMT1, DNMT3B, MTHFS, MAT2B, and GLRX were also validated in the TCGA IDH1-MT cohort **(Additional File 1, Figure S8)**.

**Figure 7.**
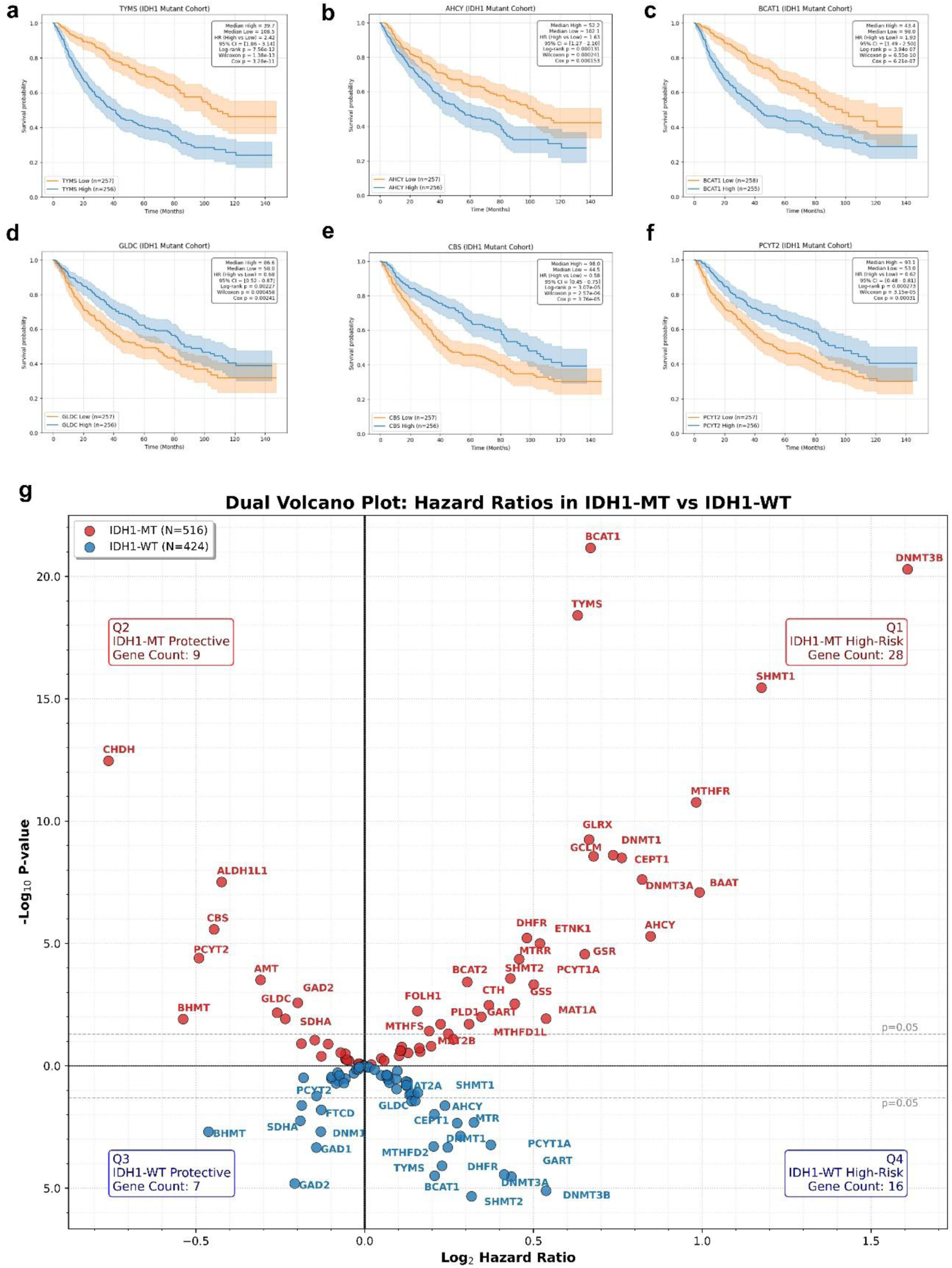
Divergent Prognostic Drivers of One-Carbon Metabolism Outline Detrimental and Protective Subtypes in IDH1-Mutant Glioma. Kaplan-Meier survival analysis illustrating prognostic outcomes within IDH1-Mutant CGGA Cohort for key drivers: **(a)** TYMS, **(b)** AHCY, **(c)** BCAT1, as detrimental proliferative drivers, where high expression is significantly associated with reduced overall survival and **(d)** GLDC, **(e)** CBS, and **(f)** PCYT2 protective drivers, where high expression confers a survival advantage. Patients were stratified into high (blue) and low (orange) expression groups. **(g)** Dual Volcano Plot comparing the hazard ratios across 64 OCM genes in IDH1-MT (red, top) versus IDH1-WT (blue, bottom) cohorts, identifying protective and high-risk drivers. The X-axis displays the Log2 Hazard Ratio (right = high risk; left = protective), and the Y-axis represents statistical significance derived using univariate Cox regression.

#### 3.5.2 Compensatory OCM Pathway Activation Confers a Protective Prognostic Advantage in IDH1-MT Tumours

The compensatory mechanisms for 1C maintenance emerged as protective prognostic genes within the IDH1-MT cohort compared to previously identified transcriptionally upregulated proliferative subset. This further validates that the high expression of specific metabolic enzymes associated with core compensatory pathways was strongly protective (HR < 1.0) in IDH1-MT tumours and that the efficiency of these compensatory mechanisms is vital for maintaining the indolent phenotype. The most protective genes, ALDH1L1 (HR=0.52, p=4.35e-07), CBS (HR=0.58, p=3.76e-05), PCYT2 (HR=0.62, p=0.00031), BHMT (HR=0.68, p=0.00258), and GLDC (HR=0.68, p=0.00241), are fundamentally involved in managing 1C unit generation and lipid-membrane homeostasis **(Figure 7D-F)**. The other low risk genes conferring protective advantage include GAD2 (HR=0.71, p=0.00704), CHDH (HR=0.44, p=6.54e-10), and AMT (HR=0.69, p=0.00459). The protective effects of CBS (HR=0.56, p=0.018), GLDC (HR=0.59, p=0.0277), BHMT (HR=0.57, p=0.0176), and PCYT2 (HR=0.41, p=0.000474) were also validated in the TCGA IDH1-MT cohort **(Additional File 1, Figure S9)**. High expression of GLDC, AMT, and AHCYL2 (HR=0.6, p=0.0348), a regulatory pseudo enzyme known to slow down AHCY function and key components of the mitochondrial 1C generating axis, suggests a strong capacity to feed the OCM cycle is necessary to sustain methylation demands without driving excessive *de novo* synthesis pathways, further preserving the low-grade metabolic profile characteristic of IDH1-MT tumours.

#### 3.5.3 Elevated OCM Activity is Universally Detrimental in Maximally Aggressive IDH1-WT Tumours

The individual gene prognosis generally mirrored the high-risk signature in the IDH1-WT, further strengthening the notion that high OCM activity is inherently deleterious in this background. Thus, in contrast to the divergent outcomes in IDH1-MT disease, the prognostic genes in the IDH1-WT cohort were associated with poorer prognosis, confirming that any degree of OCM activation adversely affects survival. For example, highly aggressive drivers correlated with worse outcomes, such as TYMS (CGGA HR = 1.64, p=9.61e-06; TCGA HR = 1.41, p=0.0322), SHMT2 (CGGA HR = 1.46, p=0.000635; TCGA HR = 1.48, p=0.0133), and GLRX (TCGA HR = 1.81, p=0.000223). Other genes showing highly significant detrimental effects in the robust CGGA dataset include BCAT1 (HR=1.52, p=0.00015), DHFR (HR=1.47, p=0.00045), and DNMT3B (HR=1.52, p=0.000147), confirming that high proliferative and metabolic capacity consistently correlates with maximally aggressive clinical behaviour in IDH1-WT tumours **(Additional File 1, Figure S10A-F)**. Conversely, the metabolic dependencies imposed by the IDH1 mutation showed altered behaviour of key prognostic genes. Several heavily protective genes identified in the IDH1-MT cohort completely lost their survival benefit in the IDH1-WT environment, such as ALDH1L1 (HR = 0.93, p = 0.522) and CBS (HR = 1.15, p = 0.196) **(Additional File 1, Figure S10G-H)**. Most strikingly, GLDC demonstrated a complete reversal of its biological role, while being highly protective in IDH1-MT tumors (HR = 0.68, p = 0.00241), it became a significant detrimental risk factor **(Additional File 1, Figure S10I)** in the CGGA IDH1-WT cohort (HR = 1.37, p = 0.00483). Genes that were effectively suppressed or non-significant in the mutant cohort emerged as significant detrimental drivers in the IDH1-WT, including MTHFD2 (HR = 1.25, p = 0.0458), MAT2A (HR = 1.40, p = 0.00257), and PCYT1A (HR = 1.41, p = 0.00196) **(Additional File 1, Figure S10J-L).**

Taken together, the results show that in aggressive pan-glioma and IDH1-WT cohorts, high expression of OCM genes is associated with poor prognosis, reflecting the high metabolic demands of these tumours. In both the CGGA and TCGA datasets, the aggressive pan-glioma and IDH1-WT cohorts are defined by the presence of detrimental OCM genes. However, within the generally indolent IDH1-MT cohort, the individual prognostic analysis across both CGGA and TCGA supports the view that the type of OCM activity defines the prognostic outcome in IDH1-MT gliomas. Although the combined score remains strongly detrimental in IDH1-MT tumours, this indicates that the weight and magnitude of the detrimental metabolic shifts and catabolic pathways confirmed across datasets may override the beneficial effects and act like metabolic aggressors, providing an advantage of the protective compensatory axes, thereby acting as drivers of poor survival in this subset. MTHFS and PCYT2 were conserved across all datasets and universally detrimental and protective, respectively. CBS, GLDC, BHMT, and PCYT2 were universally protective in IDH1-MT **(Table 2)**. This differential regulation and opposing prognostic roles of individual genes suggest that activation of the OCM pathway is a major factor driving prognostic heterogeneity in IDH1-MT gliomas. Targeting the aberrant activation of these proliferative OCM genes in high-risk IDH1-MT tumours represents a promising therapeutic approach. The summary of prognosis across 64 OCM genes between IDH1-WT and IDH1-MT in the CGGA cohort is presented in a dual volcano plot **(Figure 7G)**.

**Table 2:** Comprehensive summary of One-Carbon Metabolism gene prognosis across TCGA and CGGA glioma cohorts. Total significant prognostic genes are identified using a Cox proportional hazards regression with a p-value < 0.05. Prognostic directions are categorised into detrimental drivers (Hazard Ratio [HR] > 1.0), in which elevated expression correlates with poorer overall survival and greater disease aggressiveness, and protective drivers (HR < 1.0), which confer a survival advantage.

| Cohort | Total Significant Prognostic Genes (Cox p < 0.05) | Detrimental (HR > 1.0, Cox p < 0.05) | Protective (HR < 1.0, Cox p < 0.05) |
| --- | --- | --- | --- |
| IDH1-WT | CGGA<br>N = 25 | n=18, AHCY, BCAT1, CHPT1, DHFR, DNMT1, DNMT3A, DNMT3B, GART, GLDC, MAT2A, MTHFD1, MTHFD2, MTHFS, MTR, PCYT1A, SHMT2, TCN2, TYMS | n=7, BHMT, DMGDH, GAD1, GAD2, PCYT2, SDHA, SOD1 |
|  | TCGA<br>N = 9 | n=7, AHCY, GLRX, GSS, MTFMT, MTHFS, SHMT2, TYMS | n=2, PCYT1B, PCYT2 |
| IDH1-MT | CGGA<br>N = 31 | n=23, AHCY, BAAT, BCAT1, BCAT2, CEPT1, DHFR, DNMT1, DNMT3A, DNMT3B, ETNK1, GART, GCLM, GLRX, GSR, GSS, MAT2B, MTHFD1L, MTHFR, MTHFS, MTRR, SHMT1, SHMT2, TYMS | n=8, ALDH1L1, AMT, BHMT, CBS, CHDH, GAD2, GLDC, PCYT2 |
|  | TCGA<br>N = 16 | n=10, CEPT1, DNMT3A, DNMT3B, ETNK1, GLRX, GSR, MAT2B, MTHFS, MTRR, SHMT1 | n=6, AHCYL2, BHMT, CBS, GLDC, PCYT2, SDHA |

### 2.6 IDH1 Mutation Status Dictates OCM-Driven Remodelling of the Tumour Immune Microenvironment

The tumour immune microenvironment (TIME) analysis of the OCM gene signature revealed IDH1 mutation status-dependent differential regulation and a strong metabolic association. Correlation analysis of immune associations between the OCM gene signature and 22 immune cell types revealed significant differences between the IDH1-WT and IDH1-MT cohorts. In the IDH1-MT, M2-macrophages exhibited the strongest correlation with the overall signature score (R = −0.227, P = 3.16E-06) **(Figure 8A, Additional File 2, Table S5)**. This inverse relationship suggests that a highly active 1C metabolic state, corresponding to a low signature score, could drive a reduction in M2-polarised macrophages. Other immune infiltrates showing significant negative correlation included dendritic cells resting (R = −0.162, P = 0.00095), T cells CD4 memory activated (R = −0.127, P = 0.0096) and T cells gamma delta (R = −0.101, P = 0.0407). In contrast, monocytes alone displayed the highest positive correlation (R = 0.185, P = 0.00015) in the IDH1-MT tumours, further suggesting differential regulation of specific myeloid cell maturation and recruitment pathways, thereby promoting monocyte infiltration while simultaneously repressing M2 polarisation. However, the IDH1-WT cohort showed a considerable shift in the correlation profile structure **(Figure 8B)**. Activated NK cells showed the most significant negative correlation (R = −0.156, P = 0.0075), followed by resting NK cells, with the strongest positive correlation (R = 0.139, P = 0.0178). Macrophages M1 and Neutrophils also displayed significant positive correlations in IDH1-WT tumours (R = 0.120, P = 0.041, and R = 0.116, P = 0.048, respectively). Importantly, the M2-macrophages did not display a significant correlation in IDH1-WT tumours (R=0.106, P=0.071) **(Additional File 2, Table S6)**.

**Figure 8.**
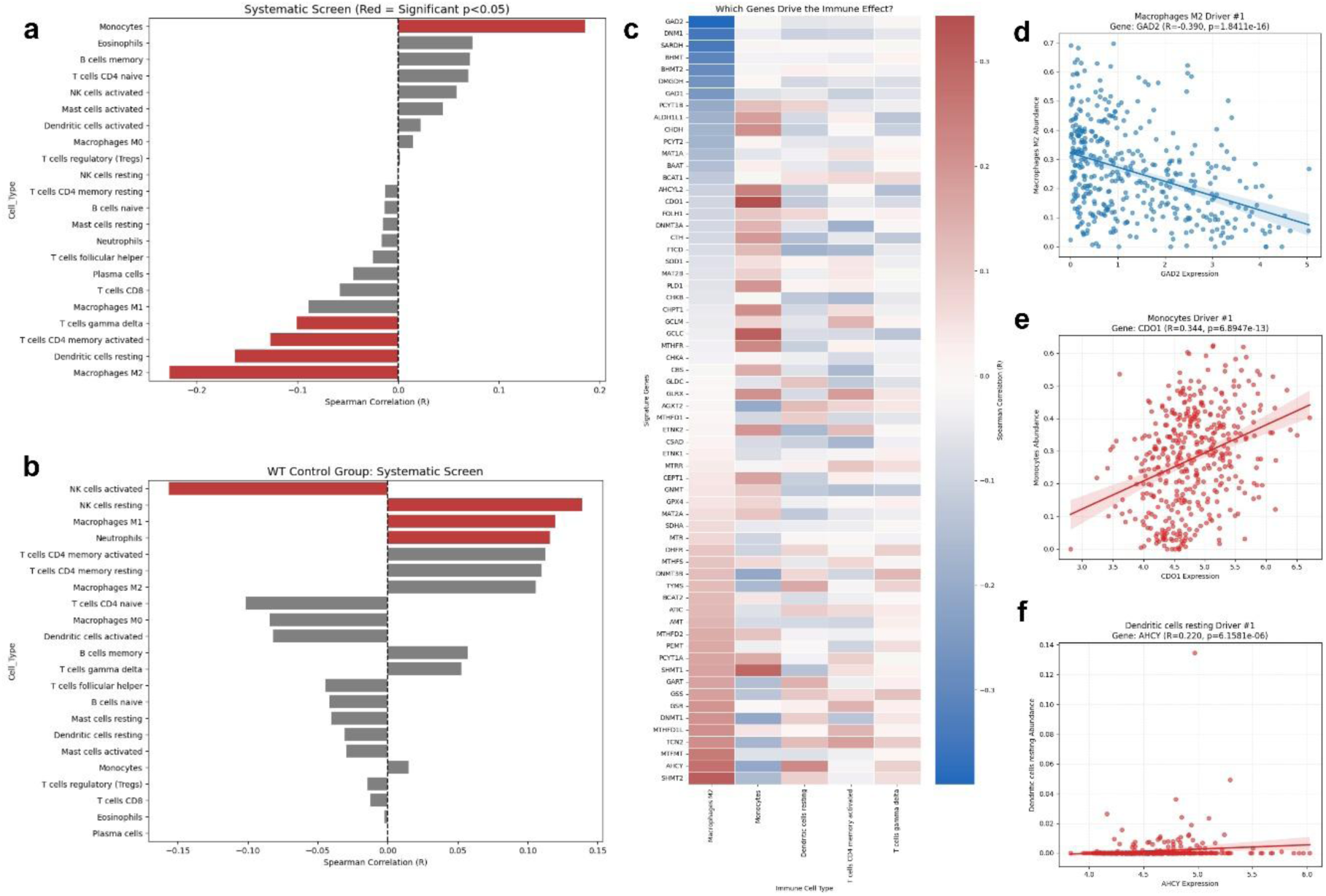
The OCM Signature Selectively Remodels Myeloid Compartment Driving Monocyte Infiltration and Suppressed M2 Macrophages in IDH1-MT Gliomas. Systematic correlation screen visualising Spearman correlation (R) values across 22 immune cell types (red bars denote p < 0.05) in **(a)** the IDH1-MT highlighting specific remodelling of the myeloid lineage with significant positive correlation with monocytes and a strong negative correlation with immunosuppressive M2 macrophages, and **(b)** the IDH1-WT cohort showing different immunomodulatory profile associating negatively with activated NK cells rather than the myeloid compartment. **(c)** Heatmap showing 64 OCM gene correlations (Blue indicates negative correlation; red indicates positive correlation) in IDH1-MT tumours to identify immune-effect gene drivers across the top five significantly modulated immune populations. Scatter plots with linear regression (and 95% CI) displaying the top metabolic drivers for targeted immune cell abundance in the IDH1-mutant cohort: **(d)** Macrophages M2 Driver (GAD2), **(e)** Monocytes Driver (CDO1), and **(f)** Dendritic cells resting Driver (AHCY).

To further dissect OCM-immune associations in IDH1-MT, we analysed the 64 OCM gene network with the Top 5 most correlated immune cell types to identify the Top 3 OCM metabolic drivers within each group **(Figure 8C)**. In the IDH1-MT, the negative correlation with Macrophages M2 was strongly driven by genes such as GAD2 (R = −0.390, P = 1.841e-16), DNM1 (R = −0.349, P = 2.864e-13), and SARDH (R = −0.341, P = 9.793e-13) **(Figure 8D)**.

The involvement of GAD2 and DNM1 suggests that novel links between amino acid metabolism and OCM may regulate the suppressive M2 phenotype. Conversely, the monocyte correlation was positively driven by key antioxidant and folate pathway enzymes, including CDO1 (R = 0.344, P = 6.8947e-13), GCLC (R = 0.310, P = 1.2466e-10), and SHMT1 (R = 0.296, P = 8.7911e-10) **(Figure 8E)**. Although the combined signature negatively correlated with resting dendritic cells, the AHCY gene showed a positive correlation (R = 0.220, P = 6.1581e-06) **(Figure 8F)**, which was contrasted by negative correlations from FTCD and ETNK2 **(Additional File 1, Figure S11)**. This highlights that even within the IDH1-MT cohort, specific metabolic branches, such as AHCY-mediated SAM recycling, can exert distinct, cell-specific effects on the TIME composition. In IDH1-WT, the genes driving the immune associations were associated with a different subset of OCM enzymes **(Additional File 1, Figure S12)**. The strongest negative correlation among activated cytotoxic NK cells was associated with genes such as GLRX (R = −0.338, P = 3.4398e-09) and GCLM (R = −0.334, P = 5.4122e-09). In contrast, the strongest positive correlation for resting cytotoxic NK cells was associated with the genes DHFR (R = 0.214, P = 2.3884e-04) and GLRX (R = 0.2, P = 6.0701e-04).

This data collectively demonstrates that the regulatory mechanisms linking OCM to immune infiltration are distinct across IDH1 status. In IDH1-WT gliomas, high metabolic activity, enhanced by the redox regulators GLRX (R = −0.338) and GCLM (R = −0.334), correlates inversely with cytotoxic NK cell infiltration, highlighting a metabolic mechanism of immune evasion. However, in the IDH1-MT cohort, the OCM signature could act as a metabolic regulator that selectively modulates myeloid populations, thereby promoting monocyte infiltration through the CDO1, GCLC, and SHMT1 gene axis while simultaneously repressing M2 Macrophages via GAD2, DNM1, and SARDH. This functional divergence, characterised by increased monocytes and decreased M2 macrophages rather than universal suppression of immune phenotype, suggests a clear metabolic-immune axis defined by a high OCM score.

This characterisation of OCM regulation, functional validation of high-risk dependency, prognostic stratification, and associated TIME plasticity provides a preliminary biological context for discussing how aberrant activation of proliferative OCM pathways serves as a central vulnerability that could override the survival benefit conferred by IDH1 mutations in gliomas.

## 3. Discussion

The discovery of IDH mutations fundamentally reshaped the molecular landscape of neuro-oncology. These cytosolic mutations are characteristic drivers of a large subset of diffuse low-grade gliomas and secondary glioblastomas. The genetic and epigenetic consequences of IDH mutations generating oncometabolite D-2HG and resulting in G-CIMP phenotype are well-established metabolic alterations known for the survival of these tumours, and remain an area of active investigation (43). Our findings present a reflective approach to complement the metabolic architecture of IDH1-MT gliomas, with a focus on comprehensive restructuring of the OCM network. Although the study needs further investigation and validation in patient samples and tumor models, these results provide a potential roadmap for metabolic precision medicine in gliomas. This study involved analysis of transcriptional regulation of 64 OCM genes across TCGA and CGGA glioma cohorts by employing a collective OCM signature score to quantify global pathway expression, validated using *in-vitro* cell line models and nutrient deprivation assay. The prognostic significance was determined using univariate and bivariate Cox regression models, which strongly implicated OCM dysregulation as a key determinant of survival in glioma patients. Further, the functional consequences of tumour immune cell infiltration were assessed using CIBERSORTx and Spearman correlation analysis between the OCM signature and 22 immune cell infiltration profiles. Together, this study suggests that the metabolic rewiring of OCM may have prognostic and therapeutic potential.

The transcriptomic analyses revealed that IDH1-MT gliomas uniformly downregulate many OCM genes that otherwise support proliferation. This is also consistent with the well-established observation that IDH1-MT slows glioma growth and confers a survival advantage. *In vitro* and *in vivo* studies have previously shown that mutant IDH1 expression reduces TCA-cycle flux, thereby lowering NADPH/NAD+ levels and significantly inhibiting glioma cell proliferation (44). Accordingly, IDH1-MT gliomas exhibited a suppressed metabolic signature. This OCM suppression also aligns with the concept that IDH1 mutations enforce an epigenetic reprogramming towards differentiation, unlike IDH1-WT aggressive metabolic tumours (45). Our study also presents a transcriptionally repressed environment in IDH1-MT gliomas, characterised by reduced expression of proliferation-associated enzymes involved in *de novo* serine synthesis and components of the cytosolic folate cycle, alongside significant expression of TYMS, DHFR, and BCAT1 from the 1C network.

Despite the overall downregulation of OCM, we identified a distinct subset of OCM genes (such as TYMS, SHMT1, and DNMT3B) that, when elevated in IDH1-MT gliomas, are known to drive glioma aggressiveness and indicate a poor prognosis. The *in vitro* functional validation in the aggressive U87 genetic background with IDH1-MT using RT-qPCR also directly supported the existence of this high-risk metabolic profile similar to that of secondary gliomas. While basal redox pathways (GLRX) remained suppressed and compensatory transsulfuration (CBS) was activated, they concurrently maintained high expression of TYMS, SHMT1, and DNMT3B. This validates that within a highly aggressive genetic background, IDH1-MT cells can hijack proliferative OCM pathways to sustain rapid growth, with a poor-prognosis patient subset identified earlier. These genes encode rate-limiting enzymes in nucleotide and methyl-donor biosynthesis, and their high expression is known to drive glioma aggressiveness. Previous studies have shown that overexpressed SHMT1 (a 1C donor from serine) in low-grade gliomas correlates with shorter patient survival, along with TYMS and DHFR. Reports suggest that SHMT1 overexpression activates mTORC1 signalling, thereby driving proliferation and invasion and leading to increased growth in indolent tumours (46,47). Oncogenic methyltransferase (DNMT3B) was likewise upregulated in high-grade gliomas, and its overexpression correlates with genome-wide hypermethylation and poor prognosis (48). The suppression of TYMS, DHFR, GART, and SHMT1 reflects slow growth and active metabolic necessity. IDH1-MT cells may reduce competition for limited NADPH reserves by prioritising redox maintenance over DNA synthesis. This is further supported by the concurrent downregulation of other NADPH-dependent lipid synthesis genes such as CHPT1 and CEPT1. Clinically, IDH1-MT patients that show this high-OCM signature effectively resemble the prognosis of IDH1-WT gliomas, further challenging the conventional notion of upregulated biosynthetic fluxes in cancer metabolism (49). Therefore, our study indicates that IDH1-MT gliomas exhibit a two-state metabolic phenotype: a basal state of metabolic quiescence characterised by downregulation of high-demand pathways, and a stress-responsive state characterised by upregulation of compensatory 1C and transsulfuration enzymes. A subset-specific upregulation of these proliferation-promoting genes contributes to an aggressive phenotype. Mechanistically, this might reflect the complex epigenetic state induced by IDH1 mutations. Another interesting finding showed downregulation of *de novo* DNA methyltransferases (DNMT3A, DNMT3B) and SAM synthase (MAT1A) in IDH1-MT tumours. The global hypermethylation (G-CIMP) phenotype in IDH1-MT tumours is thus primarily driven by D-2HG inhibiting TET-family demethylases, rather than by increased methyltransferase activity, as further supported by suppression of AHCY, which leads to SAH buildup and inhibition of methyltransferases.

In contrast, we observed that some mitochondrial OCM enzymes, such as MTHFD2, GLDC, and GCLC, are upregulated in IDH1-MT tumours, and they correlate with better survival outcomes. In many cancers, MTHFD2 is considered pro-tumorigenic (50,51), but a recent study by Huang et al. (52) found that lower MTHFD2 expression in high-grade tumours predicts poor prognosis. In contrast, higher expression in low-grade gliomas is significantly associated with improved patient survival, consistent with our findings. They showed that MTHFD2 overexpression in glioma cells inhibited migration, invasion, and proliferation by suppressing the ERK pathway (52). The upregulation of GLDC in our patient data suggests that IDH1-MT tumours preferentially utilise glycine catabolism to fuel the 1C pool, rather than the serine synthesis pathway (PHGDH) that is often upregulated in WT tumours (53). This explains that IDH1-MT tumours, particularly the 1p/19q co-deleted subset (which often lose PHGDH due to 1p deletion), are hypersensitive to glycine restriction or GLDC inhibition (54). The *in-vitro* glycine serine starvation assay also established the critical need for upregulated GLDC for IDH-MT tumours. While the wildtype cells retained the flexibility to upregulate the serine biosynthesis axis and survive the nutrient deprivation, IDH1-MT cells exhibited a strict functional auxotrophy. The MT cells were unable to achieve metabolic rescue under serine or glycine restriction, resulting in a lethal population collapse (21.58% viability) during synergistic dual deprivation over 96 hours. Further, GCLC (glutathione synthesis subunit) was elevated in the IDH1-MT high-OCM subset, supporting redox defence, which is beneficial in the oxidative tumour environment of IDH1-MT. Studies show glioblastoma cells overexpress GCLC downstream of telomerase (via FOXO1) to promote proliferation (55). This study also complements prior findings by suggesting that the elevated GCLC subgroup in IDH1-MT scored better, with enhanced glutathione metabolism partly counteracting oxidative stress in these tumours. Taken together, the compensatory upregulation of mitochondrial folate/glycine catabolism and glutathione production in IDH1-MT gliomas correlates with a less aggressive phenotype, highlighting conditional, dependent roles of these pathways that reinforce the notion that the canonical IDH1-MT metabolic state relies on mitochondrial support and that transsulfuration is less aggressive than cytosolic proliferation. Therefore, computational and functional assays in our study complement these findings and bridge them, demonstrating that even when IDH1-MT tumours activate the proliferative OCM machinery, they remain intricately dependent on IC sources and compensatory salvage pathways to drive aggressive growth.

Further, we investigated the OCM transcriptional landscape of IDH1-MT astrocytomas altered by canonical TP53 and ATRX co-mutations. Tombari et al. showed that gain-of-function TP53 mutants enhance serine/glycine synthesis and uptake of essential amino acids. In TP53-MT breast cancer cells, silencing TP53 reduced labelled glucose flux into serine/glycine (56). Concurrent with this, our results show similar activation of serine/glycine pathway genes (such as MTHFR, SHMT1/2, MTHFD2) to compensate for IDH1-MT metabolic stress, while wildtype TP53 restricts anabolic metabolism. We also found that high OCM signature scores correlated with a higher frequency of ATRX mutations in the IDH1-MT cohort. Previous studies by Hariharan et al. showed that ATRX loss in glioma cells sensitises them to dsRNA agonists, leading to an increase in interferon-stimulated gene expression and T-cell infiltration, and their co-expression with IDH1-MT reduces these immune responses (57). We also showed these high-OCM subgroups were enriched across classic astrocytoma genotypes (IDH1 mutant with TP53/ATRX loss) and IDH1-MT oligodendrogliomas (with 1p/19q codeletion and TERT but no ATRX mutation). This suggests that while IDH1 initiates metabolic rewiring, the loss of ATRX synergises with OCM dysregulation to drive the tumour towards harmful proliferative or compensatory mitochondrial axes.

To further enhance our understanding and uncover a novel link between OCM activity and the TIME, we evaluated the 1C transcriptional state in gliomas with immune microenvironmental changes that differed by IDH status. IDH1-WT gliomas exhibited higher overall immune cell infiltration and a more immunosuppressive phenotype than IDH1-MT gliomas, consistent with the literature (58). Recent profiling studies also find that IDH-WT tumours are enriched for monocyte-derived macrophages with tumour-promoting phenotypes (59,60), but our data reveal a structured segregation mediated by OCM. We found that the OCM signature is not immune-neutral but actively alters the composition of the myeloid compartment. We observed a negative correlation between the OCM signature and M2 macrophages (driven by GAD2 and DNM1), and a positive correlation with monocytes (driven by CDO1 and GCLC) in IDH1-MT tumours. This suggests that the distinct metabolic milieu in IDH1-MT tumours creates an environment that restricts macrophage polarisation towards the immunosuppressive M2 phenotype. This metabolic exclusion of M2 macrophages contributes to the indolent and less aggressive immune state characteristic of IDH-MT gliomas. Our data indicate that upregulated GAD2 in low OCM-score IDH1-MT cancers, along with GABA signalling, could promote M2 macrophage polarisation and an immunosuppressive phenotype. In IDH1-MT gliomas, a GABA-driven M2-dominant microenvironment might represent a quiescent tissue state that suppresses inflammation-driven angiogenesis and invasion compared with the inflammatory, immune-invasive myeloid landscape of IDH1-WT GBM. High expression of monocyte-driving genes implies significant oxidative stress induced by chemokines such as CCL2, highlighting high ROS pressure in aggressive, high-OCM IDH1-MT tumours. In contrast, the high-redox environment of IDH1-WT tumours (high GLRX) correlates with NK cell exclusion, suggesting a metabolic mechanism of immune evasion where tumour antioxidant capacity neutralises NK-mediated cytotoxicity. These findings align with those of Bunse et al., who demonstrated that infiltrating myeloid cells remain in an immunosuppressive state in IDH-MT gliomas. In contrast, in late-stage tumours, monocyte-derived macrophages impose tolerance via the aryl hydrocarbon/tryptophan pathway (61).

Therefore, this study demonstrates that OCM is not a static background feature of IDH1-MT gliomas but a dynamic determinant of tumour behaviour. While the prognostic power of the OCM signature was robust in the CGGA and pan-glioma cohort analyses, the attenuated statistical significance in the TCGA IDH1-MT subset warrants further investigation. This suggests that underlying differences in patient demographics or treatment protocols might influence metabolic signals, dictating further validation. Additionally, while our *in vitro* study confirmed dependence on serine/glycine flux and modulation of metabolic reprogramming, future studies should employ clinical validation in high- and low-risk models, patient samples, and *in vivo* experiments. Overall, this research advances the field of neuro-oncology by moving beyond genomic classification towards metabolic precision medicine. The reliance of classical IDH1-MT tumours on CBS and GLDC creates a synthetic lethal opportunity. Inhibitors of CBS or the glycine cleavage system could trigger a redox crisis in these NADPH-depleted cells, thereby selectively killing them. This could be particularly relevant for 1p/19q co-deleted oligodendrogliomas that lack PHGDH and are strictly dependent on exogenous serine/glycine and transsulfuration. For high-risk IDH1-MT patients identified using the OCM prognostic signature, standard IDH1 inhibitors might be insufficient. These tumours may have metabolic profiles resembling GBM and potential sensitivity to antifolates. Targeting the specific metabolic vulnerabilities defined by the OCM signature offers a promising strategy to improve survival outcomes for patients with IDH1-MT gliomas.

## 4. Conclusion

This study systematically characterised the transcriptional landscape of OCM in diffuse gliomas and established it as a potential key determinant of clinical behaviour and a therapeutic target. We demonstrated that IDH1 status is the fundamental driver of OCM rewiring, which induces a global suppression of high-demand proliferative pathways and necessitates the compensatory activation of 1C sources, thereby identifying a two-state metabolic phenotype within the IDH1-MT cohort. While IDH1 mutations generally confer a survival advantage, we identified a high-risk subset of these tumours where the aberrant reactivation of proliferative OCM genes could drive an aggressive phenotype resembling GBMs. Conversely, the upregulation of compensatory genes was strongly protective and marked the indolent phenotype. Though this divergent phenotype in IDH-MT gliomas requires further investigation and validation, it suggests that the OCM signature can be used for risk stratification and as a therapeutic vulnerability. Furthermore, our results show a metabolic-immune axis governed by the OCM signature that acts as a metabolic regulator in TIME. High metabolic flux was associated with oxidative stress-driven monocyte recruitment. At the same time, the metabolically quiescent state favoured M2 macrophage polarisation, suggesting that metabolic interventions could potentially remodel the immune landscape in IDH1-MT gliomas, offering new avenues for combination therapies. Thus, our study complements existing work by providing a framework for resolving the OCM landscape into protective and aggressive signatures for the clinical heterogeneity of IDH1-MT tumours and helps categorise actionable metabolic targets that could redefine the therapeutic standard of care.

## 5. Materials and Methods

### 5.1 Data collection and preparation

The dataset for this study was constructed using omics data from two independent, large-scale datasets, The Cancer Genome Atlas (36) (TCGA.) and The Chinese Glioma Genome Atlas (37) (CGGA). To facilitate cross-cohort comparability and minimise batch effects from raw data processing, we used the GlioVis (38) data portal to retrieve expression matrices and clinical metadata. In accordance with the WHO 2021 Classification of Tumours of the Central Nervous System (39), which emphasises molecular diagnosis over histological grading, we used the merged GBMLGG cohort available through GlioVis in TCGA. This merged dataset aggregates samples from the TCGA-LGG (Lower-Grade Glioma, WHO Grades II and III) and TCGA-GBM (Glioblastoma, WHO Grade IV) projects into a unified matrix. We retrieved the normalised mRNA expression datasets, ensuring that transcript abundance was comparable across samples with varying sequencing depths. We also extracted the corresponding clinical phenotypes directly from GlioVis to ensure perfect sample matching with variables including Overall Survival (OS) Time, Censoring Status, IDH1 Mutation Status, and 1p/19q Co-deletion Status. These served as the primary stratification variable in the dataset, filtered by defined IDH1 status (Wildtype vs. Mutant), and, to further refine the molecular classification, distinguishing between IDH-MT astrocytomas (non-codel) and oligodendrogliomas (codel). The validity of the identified metabolic signatures was also assessed in an independent CGGA cohort, accessed via the GlioVis software to maintain consistency in data formatting and preprocessing.

The OCM gene network defined the biological scope of the study. We retrieved the OCM gene set defined by the WikiPathways Database (40), utilising pathway identifier WP2525 (One Carbon Metabolism, Transsulfuration and Related Pathways). This pathway map provided a comprehensive curation of the enzymes and transporters involved in the folate cycle, methionine cycle, and the transsulfuration pathway. A total of 64 candidate genes were extracted for analysis **(Additional File 2, Table S1)**.

### 5.2 Statistical thresholding, signature score and Cohen’s d calculation

While individual gene analysis provides specific insights, quantifying the aggregate activity of the OCM pathway was necessary for prognostic modelling. A composite signature score and Cohen’s d were used to quantify the magnitude of separation between biological groups. Z-score normalisation was used to compare gene expression levels, enabling calculation of a composite signature score. Raw normalised expression values (FPKM or TPM) were standardised gene-wise using equation (1) within each cohort. For a given gene (g) and patient (i) in cohort, the Z-score (Zg.i) was calculated as:

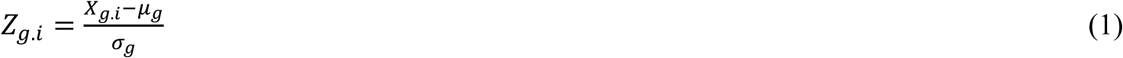

Where, *X_g_*_.*i*_ is the raw expression value of gene (g) for patient (i), *μ*g is the mean expression of gene (g) across all patients in the cohort, and *σ_g_* is the standard deviation of gene (g) across all patients in the cohort. This transformation centres the expression distribution of every gene at zero with a standard deviation of one, which was critical for equal weighting and cohort independence. This ensured that genes with high baseline expression didn’t disproportionately influence the combined score compared to low-abundance regulatory enzymes, while maintaining the internal distributional properties of each dataset.

The combined signature score was calculated for each patient using equation (2) to summarise the metabolic risk profile into a single continuous variable. This score represented the arithmetic mean of the Z-scored expression values of the genes comprising the signature and was calculated as:

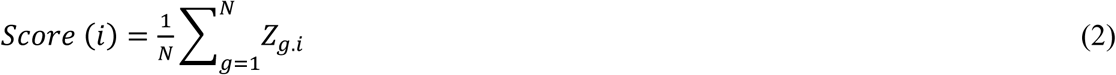

Where N is the number of genes in the signature, and Zg.i is the Z-score of gene (g) for patient (i). This additive model assumes that the simultaneous upregulation of multiple genes in the deleterious metabolic state cumulatively contributes to the aggressive phenotype. Further, the raw p-values were also adjusted for multiple hypothesis testing using the Benjamini-Hochberg (BH) procedure to control the False Discovery Rate (FDR). We defined Differentially Expressed Genes (DEGs) based on a stringent statistical threshold of an FDR-adjusted p-value < 0.05.

While p-values indicate statistical significance, they do not convey the magnitude of the biological difference, especially in large datasets where small differences can become statistically significant. To rigorously quantify the magnitude of the transcriptional shift between IDH1-WT and IDH1-MT tumours, we calculated Cohen’s d for each gene in the signature using equations (3) and (4). Cohen’s d was computed as the difference between the means of the two groups divided by the pooled standard deviation.

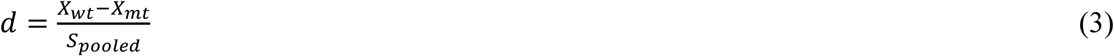

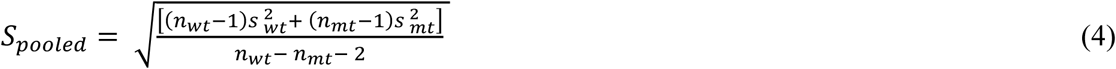

This metric is used in the datasets where even small differences become statistically significant. The effect size was interpreted as: (a) Positive d: Indicated genes that were upregulated in IDH1-WT tumours (and consequently suppressed in IDH1-MT). A large positive d suggests the gene is part of the proliferative axis driven by the wildtype phenotype, and (b) Negative d: Indicated genes that were upregulated in IDH1-MT tumours (and consequently suppressed in IDH1-WT). These genes represent the compensatory metabolic adaptations required to survive the stress of the IDH1 mutation. This metric allowed us to rank genes not only by their p-values but also by their contribution to the biological separation of the two tumour entities, thereby helping identify true drivers of the metabolic shift.

### 5.3 Transcriptomics expression analysis

Following data acquisition, the computational pipeline for expression plots was implemented to process and analyse the transcriptomic data. To address the integrated multi-cohort data, we first harmonised the sample identifiers. TCGA barcodes (e.g., TCGA-XX-YYYY-01A-ZZZ) were construed to extract the patient-level identifier (the first 12 characters, e.g., TCGA-XX-YYYY) or the sample-level identifier (first 15 characters). Such truncation was done in order to make the expression data consistent with the mutation and clinical data. In cases where multiple data points were associated with the same patient, an aggregation function was used to reduce the duplicates, such that each patient was represented by one expression vector in order to prevent sample size inflation and violations of statistical testing assumptions.

We used the heatmap R package and the seaborn library in Python to visualise OCM global gene expression architecture utilising hierarchical clustering (Euclidean distance, Ward’s linkage) to group patients and genes corresponding to the IDH-WT and IDH-MT phenotypes. Additionally, volcano plots were built to highlight 64 OCM signature genes against the background transcriptome and graphically show the correlation between statistical significance (-log10 FDR) and effect magnitude (logFC).

### 5.4 Principal component analysis

PCA was used to assess the ability of the 64-gene OCM signature to distinguish between biological subtypes. It served as a dimensionality-reduction technique, transforming the high-dimensional gene expression space into a set of principal components that explain the maximum variance in the data. The analysis was performed using the PCA class from the scikit *decomposition* module in Python. The input matrix consisted of Z-score-normalised expression values for the 64 OCM genes across all samples. The parameter scale = TRUE was set to enforce unit variance, preventing genes with high variance from skewing the components. The first two principal components (PC1 and PC2) were extracted and plotted as a 2D scatter plot. Samples were colour-coded by IDH1 status (WT vs. MT) to visualise the separation.

### 5.5 Driver gene analysis

The somatic mutation data for IDH1, TP53, ATRX, and TERT in adult diffuse gliomas were obtained from TCGA whole-exome sequencing mutation calls through the UCSC Xena Browser. Sample identifiers were truncated to the first 12 characters of the TCGA barcode to ensure patient-level matching across expression and mutation datasets, as explained earlier. Gene identifiers were curated by mapping Ensembl gene IDs to official gene symbols. The Z-Score Normalisation and Signature Score Calculation was computed as explained earlier. Somatic mutation data were parsed to identify samples harbouring at least one non-silent mutation in IDH1, TP53, ATRX, or TERT. For each gene, samples were classified as: mutated (1): ≥1 somatic mutation detected, and non-mutated (0): no mutation detected. Mutation status was encoded as binary variables (IDH1_mut, TP53_mut, ATRX_mut, TERT_mut) and merged with the signature score table to generate an aligned analysis matrix. No filtering was performed based on mutation type, variant allele frequency, or functional consequence, consistent with established TCGA mutation-enrichment analyses. To identify genes within the 64-gene signature that most strongly contributed to mutation-associated expression shifts, driver gene analyses were conducted separately for IDH1, TP53, and ATRX. For each gene in the signature, the samples were divided into mutated and non-mutated groups based on each mutation, and differences in expression were assessed using the Mann–Whitney U test. The effect sizes were computed using Cohen’s d, and directionality of change was quantified as the difference in mean Z-score (Δ = WT − MT), and (Δ = MT − WT) for both IDH1 and driver mutations (TP53, and ATRX) respectively. Multiple hypothesis testing correction was performed using the FDR procedure across all 64 genes. Genes were ranked by statistical significance and effect size, and the top-ranked genes were designated as mutation-associated drivers of the signature shift. For each mutation, the volcano plots were generated using effect size on the x-axis and −log10(p-value) on the y-axis. The top-20 driver gene bar plots were generated using Cohen’s d, ordered by magnitude and coloured with horizontal significance thresholds corresponding to p-value ≤ 0.05.

### 5.6 Identification of IDH1-Specific Signature Drivers

Samples were divided into high and low signature groups based on the median signature score. Enrichment of TP53, ATRX, and TERT mutations within high-signature samples was assessed using Fisher’s exact test, with contingency tables constructed from mutation status and signature group. Odds ratios were computed using the Haldane–Anscombe correction to account for zero-cell counts. Chi-square tests were additionally reported where assumptions were satisfied. To determine whether the effects of TP53 and ATRX were independent of IDH1 status, all enrichment and score-comparison analyses were repeated separately within the IDH1-WT and IDH1-MT subsets. This stratified approach enabled the dissection of mutation-specific versus lineage-dependent effects. A multivariable logistic regression model was constructed to assess the independent contribution of TP53, ATRX, and TERT mutations to high signature activity. The dependent variable was a high signature score (binary), while predictors included binary mutation status variables and IDH1 mutation status as a covariate. Model coefficients were expanded to obtain odds ratios and reporting 95% confidence intervals with Wald test p-values. Model fitting was performed using maximum likelihood estimation. To identify signature genes uniquely associated with IDH1 mutation biology, genes significantly associated with IDH1 mutation (FDR < 0.05) were compared against TP53- and ATRX-associated driver lists using set-based analysis. A Venn diagram was generated to visualise overlaps, and genes exclusive to IDH1 were extracted as candidate IDH1-specific biomarkers.

### 5.7 Survival and prognostic analysis

The survival analysis was conducted to establish the clinical relevance of the OCM signature using the survival libraries available in Python (lifelines) and R (survival, survminer). The cohort was stratified into high-score and low-score groups based on the OCM signature score. The median score served as the primary cutoff to ensure balanced group sizes. Survival probabilities over time were estimated using the Kaplan-Meier (KM) method. The statistical significance of the difference between the survival distributions was assessed using the Log-Rank (Mantel-Cox) test. This non-parametric test compared the observed number of events in each group with the expected number under the null hypothesis of equivalent survival. To quantify the risk associated with the OCM signature, univariate Cox proportional hazards regression model was used using the *coxph* function. The signature score was modelled both as a continuous variable (to assess linear risk scaling) and as a categorical variable (high vs. low). The model output the hazard ratio (HR), which represented the relative risk of death for the high-OCM group compared to the low-OCM group (or per unit increase in score). An HR > 1 indicates that higher metabolic activity is detrimental, whereas an HR < 1 indicates a protective effect. The p-value ≤ 0.05 were considered statistically significant. The analysis was performed systematically across three conditions for both TCGA and CGGA cohorts: a) Pan-Glioma: assessing the signature’s power across the entire dataset, b) IDH-WT: assessing stratification within the aggressive, GBM-like subset, and c) IDH-MT: assessing the ability to distinguish the aggressive subset within the generally favourable mutant group.

### 5.8 Bivariate Cox regression analysis

Given that IDH1 mutation status is the single most significant prognostic factor in gliomas, we employed bivariate Cox proportional hazards models to assess whether the prognostic value of the OCM signature is independent of IDH1 mutation status. The primary goal was to assess whether the Signature Score remained statistically significant (p < 0.05) after adjustment for IDH1 status. The bivariate models incorporated two simultaneous covariates: a) Combined Score Z Filtered (X_1), included as a continuous variable to preserve the full resolution of the risk score, and b) IDH Status (X_2), encoded as a binary variable (Wildtype = 1, Mutant = 0). The p-value (≤ 0.05) was used to confirm OCM signature is clinically significant in prognosis even after controlling for the confounding factor of IDH1. To determine the predictive capability of the model, concordance index (C-index) value was calculated. This represents the probability that for a randomly selected pair of patients, the individual with the high-risk score will have a shorter survival time compared to the one with the low-risk score. C-index of 0.5 indicates random prediction, while 1.0 indicates perfect segregation, providing a quantitative benchmark of clinical efficacy.

### 5.9 Tumour microenvironment analysis

Recognising that tumour metabolism shapes the immune landscape, we next investigated the correlation between the OCM signature and the immune microenvironment. We utilised CIBERSORTx (Cell-type Identification by Estimating Relative Subsets of RNA Transcripts) deconvolution method to estimate the relative fractions of specific cell types based on the LM22 reference signature matrix that defined 22 distinct human hematopoietic cell phenotypes, including naive/memory B cells, plasma cells, 7 T-cell types, NK cells, and myeloid subsets. The analysis focused on identifying immune cell populations whose abundance was altered by the tumour’s metabolic state. The analysis was performed using Spearman’s rank correlation between the combined OCM signature score and the estimated relative abundance of each of the 22 immune cell types. The Spearman method was selected for its robustness to outliers and its ability to detect non-linear monotonic relationships, which are common in biological data. Correlations were computed separately for the IDH1-WT and IDH1-MT cohorts. The infiltration profiles of high-OCM vs. low-OCM were compared using Mann-Whitney U tests. These stratifications were critical for determining whether the metabolic signature exerted differential immune-modulatory effects depending on the underlying driver mutation. The coefficients (R) and significance values (p ≤ 0.05) were used to identify key associations.

### 5.10 Cell Line Models

The U87-MG glioblastoma cell line (ATCC HTB-14) and the U87 IDH1 mutant cell line (ATCC HTB-14 IG) harbouring the canonical R132H mutation were purchased from the American Type Culture Collection (ATCC) and used as cell line models. Cells were maintained in Eagle’s Modified Essential Medium (EMEM) containing 10% heat inactivated fetal bovine serum (FBS) supplemented with penicillin (100 U/mL) and streptomycin (100 mg/mL). These cells were cultured in accordance with the suppliers’ instructions. Cultures were maintained in a humidified atmosphere with 5% CO2 at 37°C and passaged every 24-48 hours.

### 5.11 Serine and glycine starvation assay

To evaluate dependence on exogenous 1C units, U87 IDH1-WT and IDH1-MT cells were cultured in parallel under two conditions: standard media containing serine and glycine, and selective media deprived of serine and glycine. The glioma cells were harvested using 0.25% Trypsin-EDTA and resuspended in deprived media. The cells were seeded into 96-well flat-bottom tissue culture plates at a density of 6 × 10³ cells per well in 100 µL of deprived culture medium. Following a 12-hour incubation, the cells were treated with 100 µL of fresh serine/glycine standard medium and selective medium containing serine or glycine at final concentrations of 10.5 mg/L and 7.5 mg/L, respectively. Cell proliferation and growth rates were monitored during continuous exposure periods of 24-96 hours. The standard and selective media were replenished every 24 hrs to remove their confounding effects. The differential reduction in growth rate between IDH1-WT and IDH1-MT cells under both control and deprived conditions was analysed using the colourimetric MTT cell viability assay. The spent media was aspirated after completion of treatment, and 100 µL of MTT reagent (0.5 mg/mL) was added to each well. The plates were then incubated in the dark at 37°C for 5-6 hours. To solubilise the formed formazan crystals, 100 µL of 100% (v/v) DMSO was added to each well and incubated for 15 mins at 37°C. Following crystal dissolution, the microplate was read using spectrophotometer at 570 nm. Two independent sets were evaluated in triplicate to calculate cell viability percentage.

### 5.12 Real-time qPCR

To confirm that the transcriptional rewiring observed in the TCGA and CGGA cohorts was biologically conserved in our cell models, we performed real-time quantitative PCR (RT-qPCR). Total RNA from IDH1-WT and IDH1-MT cells was harvested using Trizol according to the supplier’s kit protocol. RNA concentration and purity (A260/A280 ratio) were verified using spectrophotometry. The SuperscriptTM III first-strand synthesis kit (Takara, Cat. No. #6110A) was used to synthesise cDNA. Primers were designed based on gene sequences available in the NLM nucleotide database and the key OCM metabolic network, consisting of the folate cycle, methionine cycle, and transsulfuration pathway. Real-time studies were performed using an ABI Prism H7500 fast thermal cycler (Applied Biosystems, CA, USA). Each sample was run in triplicate to a final volume of 10 μl containing 1 μl of template (1:5 dilution), 10 pmol of each primer and 5 μl of Power SYBR Green PCR master mix (Applied Biosystems). Relative expression levels were normalised to housekeeping control (β-actin) in IDH1-WT vs IDH1-MT cells. Statistical significance was evaluated using a one-sample t-test comparing the normalised fold change of the IDH1-MT against a WT baseline. The data are presented as mean ± SEM of three independent replicates.

## Declarations

## Ethics approval and consent to participate

The study does not involve the use of any animal or human data or tissue other than public TCGA and CGGA patient datasets.

## Data Availability

The clinical cohort datasets analysed during the current study are publicly available in the TCGA (cancer.gov) and the CGGA repository (cgga.org.cn). The computational pipeline used in the current study is made publicly available in a GitHub repository at: https://github.com/mayankbajaj17/OCM-Divergence. The underlying raw data and analysed datasets are available under GitHub Release v1.0-Raw-Data-OCM-Divergence. All other supporting data are included in the manuscript and as additional files. The supplementary data consists of two additional files: Additional File 1 contains Supplementary Figures S1–S12, and Additional File 2 contains Supplementary Tables S1–S6.

## Author Contributions

**M.B.:** Writing original manuscript draft, visualisation, validation, investigation of *in-silico* and *in-vitro* methodology, formal analysis, and data curation. **R.K.:** Conceptualisation, writing, review and editing manuscript, data analysis, supervision, project administration, and funding acquisition.

## Acknowledgements

Part of the images was created using BioRender.com to ensure high-quality visual representations. GraphPad Prism 8.0.2 was used for statistical analysis and graphs. We also thank ANRF-PAIR for providing fellowship to Mr Mayank Bajaj.

## References

1. Ngo DC, Ververis K, Tortorella SM, Karagiannis TC. Introduction to the molecular basis of cancer metabolism and the Warburg effect. Mol Biol Rep. 2015 Apr;42(4):819–23. doi:10.1007/s11033-015-3857-y PubMed PMID: 25672512.

2. Luengo A, Li Z, Gui DY, Sullivan LB, Zagorulya M, Do BT, et al. Increased demand for NAD+ relative to ATP drives aerobic glycolysis. Mol Cell. 2021 Feb 18;81(4):691–707.e6. doi:10.1016/j.molcel.2020.12.012 PubMed PMID: 33382985; PubMed Central PMCID: PMC8315838.

3. Wang Y, Stancliffe E, Fowle-Grider R, Wang R, Wang C, Schwaiger-Haber M, et al. Saturation of the mitochondrial NADH shuttles drives aerobic glycolysis in proliferating cells. Mol Cell. 2022 Sep 1;82(17):3270–3283.e9. doi:10.1016/j.molcel.2022.07.007 PubMed PMID: 35973426; PubMed Central PMCID: PMC10134440.

4. Tennant DA, Durán RV, Gottlieb E. Targeting metabolic transformation for cancer therapy. Nat Rev Cancer. 2010 Apr;10(4):267–77. doi:10.1038/nrc2817 PubMed PMID: 20300106.

5. Warburg O, Wind F, Negelein E. THE METABOLISM OF TUMORS IN THE BODY. J Gen Physiol. 1927 Mar 7;8(6):519–30. doi:10.1085/jgp.8.6.519 PubMed PMID: 19872213; PubMed Central PMCID: PMC2140820.

6. Parsons DW, Jones S, Zhang X, Lin JCH, Leary RJ, Angenendt P, et al. An integrated genomic analysis of human glioblastoma multiforme. Science. 2008 Sep 26;321(5897):1807–12. doi:10.1126/science.1164382 PubMed PMID: 18772396; PubMed Central PMCID: PMC2820389.

7. Molenaar RJ, Radivoyevitch T, Maciejewski JP, van Noorden CJF, Bleeker FE. The driver and passenger effects of isocitrate dehydrogenase 1 and 2 mutations in oncogenesis and survival prolongation. Biochim Biophys Acta. 2014 Dec;1846(2):326–41. doi:10.1016/j.bbcan.2014.05.004 PubMed PMID: 24880135.

8. Badur MG, Muthusamy T, Parker SJ, Ma S, McBrayer SK, Cordes T, et al. Oncogenic R132 IDH1 Mutations Limit NADPH for De Novo Lipogenesis through (D)2-Hydroxyglutarate Production in Fibrosarcoma Cells. Cell Rep. 2018 Nov 6;25(6):1680. doi:10.1016/j.celrep.2018.10.099 PubMed PMID: 30404018; PubMed Central PMCID: PMC7337229.

9. Struys EA, Verhoeven NM, Ten Brink HJ, Wickenhagen WV, Gibson KM, Jakobs C. Kinetic characterization of human hydroxyacid-oxoacid transhydrogenase: relevance to D-2-hydroxyglutaric and gamma-hydroxybutyric acidurias. J Inherit Metab Dis. 2005;28(6):921–30. doi:10.1007/s10545-005-0114-x PubMed PMID: 16435184.

10. Amary MF, Bacsi K, Maggiani F, Damato S, Halai D, Berisha F, et al. IDH1 and IDH2 mutations are frequent events in central chondrosarcoma and central and periosteal chondromas but not in other mesenchymal tumours. J Pathol. 2011;224(3):334–43. doi:10.1002/path.2913

11. Wang P, Wu J, Ma S, Zhang L, Yao J, Hoadley KA, et al. Oncometabolite D-2-Hydroxyglutarate Inhibits ALKBH DNA Repair Enzymes and Sensitizes IDH Mutant Cells to Alkylating Agents. Cell Rep. 2015 Dec 22;13(11):2353–61. doi:10.1016/j.celrep.2015.11.029 PubMed PMID: 26686626; PubMed Central PMCID: PMC4694633.

12. Lu Y, Kwintkiewicz J, Liu Y, Tech K, Frady LN, Su YT, et al. Chemosensitivity of IDH1-Mutated Gliomas Due to an Impairment in PARP1-Mediated DNA Repair. Cancer Res. 2017 Apr 1;77(7):1709–18. doi:10.1158/0008-5472.CAN-16-2773 PubMed PMID: 28202508; PubMed Central PMCID: PMC5380481.

13. Sulkowski PL, Oeck S, Dow J, Economos NG, Mirfakhraie L, Liu Y, et al. Oncometabolites suppress DNA repair by disrupting local chromatin signalling. Nature. 2020 Jun;582(7813):586–91. doi:10.1038/s41586-020-2363-0 PubMed PMID: 32494005; PubMed Central PMCID: PMC7319896.

14. Eckel-Passow JE, Lachance DH, Molinaro AM, Walsh KM, Decker PA, Sicotte H, et al. Glioma Groups Based on 1p/19q, IDH, and TERT Promoter Mutations in Tumors. N Engl J Med. 2015 Jun 25;372(26):2499–508. doi:10.1056/NEJMoa1407279 PubMed PMID: 26061753; PubMed Central PMCID: PMC4489704.

15. Ohka F, Ito M, Ranjit M, Senga T, Motomura A, Motomura K, et al. Quantitative metabolome analysis profiles activation of glutaminolysis in glioma with IDH1 mutation. Tumor Biol. 2014 Jun 1;35(6):5911–20. doi:10.1007/s13277-014-1784-5

16. Reitman ZJ, Jin G, Karoly ED, Spasojevic I, Yang J, Kinzler KW, et al. Profiling the effects of isocitrate dehydrogenase 1 and 2 mutations on the cellular metabolome. Proc Natl Acad Sci. 2011 Feb 22;108(8):3270–5. doi:10.1073/pnas.1019393108

17. Pollak N, Dölle C, Ziegler M. The power to reduce: pyridine nucleotides – small molecules with a multitude of functions. Biochem J. 2007 Feb 12;402(2):205–18. doi:10.1042/BJ20061638

18. Nilsson R, Jain M, Madhusudhan N, Sheppard NG, Strittmatter L, Kampf C, et al. Metabolic enzyme expression highlights a key role for MTHFD2 and the mitochondrial folate pathway in cancer. Nat Commun. 2014 Jan 23;5(1):3128. doi:10.1038/ncomms4128

19. Fawal MA, Jungas T, Davy A. Inhibition of DHFR targets the self-renewing potential of brain tumor initiating cells. Cancer Lett. 2021 Apr 10;503:129–37. doi:10.1016/j.canlet.2021.01.026

20. Choate KA, Pratt EPS, Jennings MJ, Winn RJ, Mann PB. IDH Mutations in Glioma: Molecular, Cellular, Diagnostic, and Clinical Implications. Biology. 2024 Nov;13(11):885. doi:10.3390/biology13110885

21. Biedermann J, Preussler M, Conde M, Peitzsch M, Richter S, Wiedemuth R, et al. Mutant IDH1 Differently Affects Redox State and Metabolism in Glial Cells of Normal and Tumor Origin. Cancers. 2019 Dec 16;11(12):2028. doi:10.3390/cancers11122028 PubMed PMID: 31888244; PubMed Central PMCID: PMC6966450.

22. Ducker GS, Rabinowitz JD. One-Carbon Metabolism in Health and Disease. Cell Metab. 2017 Jan 10;25(1):27–42. doi:10.1016/j.cmet.2016.08.009 PubMed PMID: 27641100; PubMed Central PMCID: PMC5353360.

23. Pianka ST, Li T, Prins TJ, Eldred BSC, Kevan BM, Liang H, et al. D-2-HG Inhibits IDH1mut Glioma Growth via FTO Inhibition and Resultant m6A Hypermethylation. Cancer Res Commun. 2024 Mar 22;4(3):876–94. doi:10.1158/2767-9764.CRC-23-0271 PubMed PMID: 38445960; PubMed Central PMCID: PMC10959073.

24. Solomou G, Finch A, Asghar A, Bardella C. Mutant IDH in Gliomas: Role in Cancer and Treatment Options. Cancers. 2023 May 23;15(11):2883. doi:10.3390/cancers15112883 PubMed PMID: 37296846; PubMed Central PMCID: PMC10252079.

25. The implications of IDH mutations for cancer development and therapy | Nature Reviews Clinical Oncology [Internet]. [cited 2023 May 31]. Available from: https://www.nature.com/articles/s41571-021-00521-0

26. Bunse L, Pusch S, Bunse T, Sahm F, Sanghvi K, Friedrich M, et al. Suppression of antitumor T cell immunity by the oncometabolite (R)-2-hydroxyglutarate. Nat Med. 2018 Aug;24(8):1192–203. doi:10.1038/s41591-018-0095-6

27. Amankulor NM, Kim Y, Arora S, Kargl J, Szulzewsky F, Hanke M, et al. Mutant IDH1 regulates the tumor-associated immune system in gliomas. Genes Dev. 2017 Apr 15;31(8):774–86. doi:10.1101/gad.294991.116 PubMed PMID: 28465358; PubMed Central PMCID: PMC5435890.

28. Zhang L, Sorensen MD, Kristensen BW, Reifenberger G, McIntyre TM, Lin F. D-2-Hydroxyglutarate Is an Intercellular Mediator in IDH-Mutant Gliomas Inhibiting Complement and T Cells. Clin Cancer Res. 2018 Nov 1;24(21):5381–91. doi:10.1158/1078-0432.CCR-17-3855

29. Valvi S, Fouladi M, Fisher MJ, Gottardo NG. IDH mutant high-grade gliomas. Front Mol Neurosci. 2025 Aug 29;18. doi:10.3389/fnmol.2025.1662414

30. Zhang Z, Shi Q, Sturgis EM, Spitz MR, Wei Q. Polymorphisms and haplotypes of serine hydroxymethyltransferase and risk of squamous cell carcinoma of the head and neck: a case–control analysis. Pharmacogenet Genomics. 2005 Aug;15(8):557. doi:10.1097/01.fpc.0000170915.19522.b2

31. Liu F, Liu Y, He C, Tao L, He X, Song H, et al. Increased MTHFD2 expression is associated with poor prognosis in breast cancer. Tumor Biol. 2014 Sep 1;35(9):8685–90. doi:10.1007/s13277-014-2111-x

32. Koseki J, Konno M, Asai A, Colvin H, Kawamoto K, Nishida N, et al. Enzymes of the one-carbon folate metabolism as anticancer targets predicted by survival rate analysis. Sci Rep. 2018 Jan 10;8(1):303. doi:10.1038/s41598-017-18456-x

33. Ducker GS, Rabinowitz JD. One-Carbon Metabolism in Health and Disease. Cell Metab. 2017 Jan 10;25(1):27–42. doi:10.1016/j.cmet.2016.08.009 PubMed PMID: 27641100; PubMed Central PMCID: PMC5353360.

34. van der Meulen M, Ramos RC, Voisin MR, Patil V, Wei Q, Singh O, et al. Differences in methylation profiles between long-term survivors and short-term survivors of IDH-wild-type glioblastoma. Neuro-Oncol Adv. 2024 Jan 1;6(1):vdae001. doi:10.1093/noajnl/vdae001

35. Hvinden IC, Cadoux-Hudson T, Schofield CJ, McCullagh JSO. Metabolic adaptations in cancers expressing isocitrate dehydrogenase mutations. Cell Rep Med. 2021 Dec 21;2(12):100469. doi:10.1016/j.xcrm.2021.100469 PubMed PMID: 35028610; PubMed Central PMCID: PMC8714851.

36. The Cancer Genome Atlas [Internet]. [cited 2025 Dec 16]. Available from: https://www.cancer.gov/tcga

37. Zhao Z, Zhang KN, Wang Q, Li G, Zeng F, Zhang Y, et al. Chinese Glioma Genome Atlas (CGGA): A Comprehensive Resource with Functional Genomic Data from Chinese Glioma Patients. Genomics Proteomics Bioinformatics. 2021 Feb 1;19(1):1–12. doi:10.1016/j.gpb.2020.10.005

38. Bowman RL, Wang Q, Carro A, Verhaak RGW, Squatrito M. GlioVis data portal for visualization and analysis of brain tumor expression datasets. Neuro-Oncol. 2017 Jan;19(1):139–41. doi:10.1093/neuonc/now247 PubMed PMID: 28031383; PubMed Central PMCID: PMC5193031.

39. de Mendonça ML, Coletti R, Gonçalves CS, Martins EP, Costa BM, Vinga S, et al. Updating TCGA glioma classification through integration of molecular data following the latest WHO guidelines. Sci Data. 2025 Jun 4;12(1):935. doi:10.1038/s41597-025-05117-2

40. Agrawal A, Balcı H, Hanspers K, Coort SL, Martens M, Slenter DN, et al. WikiPathways 2024: next generation pathway database. Nucleic Acids Res. 2024 Jan 5;52(D1):D679–89. doi:10.1093/nar/gkad960

41. Wang XW, Ciccarino P, Rossetto M, Boisselier B, Marie Y, Desestret V, et al. IDH Mutations: Genotype-Phenotype Correlation and Prognostic Impact. BioMed Res Int. 2014;2014:540236. doi:10.1155/2014/540236 PubMed PMID: 24877111; PubMed Central PMCID: PMC4022066.

42. Chi R, Yao C, Chen S, Liu Y, He Y, Zhang J, et al. Elevated BCAA Suppresses the Development and Metastasis of Breast Cancer. Front Oncol. 2022 Jun 16;12:887257. doi:10.3389/fonc.2022.887257 PubMed PMID: 35785192; PubMed Central PMCID: PMC9243538.

43. Izquierdo-Garcia JL, Viswanath P, Eriksson P, Chaumeil MM, Pieper RO, Phillips JJ, et al. Metabolic Reprogramming in Mutant IDH1 Glioma Cells. PLOS ONE. 2015 Feb 23;10(2):e0118781. doi:10.1371/journal.pone.0118781

44. Bralten LBC, Kloosterhof NK, Balvers R, Sacchetti A, Lapre L, Lamfers M, et al. IDH1 R132H decreases proliferation of glioma cell lines in vitro and in vivo. Ann Neurol. 2011 Mar;69(3):455–63. doi:10.1002/ana.22390 PubMed PMID: 21446021.

45. Biedermann J, Preussler M, Conde M, Peitzsch M, Richter S, Wiedemuth R, et al. Mutant IDH1 Differently Affects Redox State and Metabolism in Glial Cells of Normal and Tumor Origin. Cancers. 2019 Dec;11(12):2028. doi:10.3390/cancers11122028

46. Zhao M, Tan B, Dai X, Shao Y, He Q, Yang B, et al. DHFR/TYMS are positive regulators of glioma cell growth and modulate chemo-sensitivity to temozolomide. Eur J Pharmacol. 2019 Nov 15;863:172665. doi:10.1016/j.ejphar.2019.172665 PubMed PMID: 31542479.

47. Gao Y, Jing N, Teng X, Wang Y. Serine hydroxymethyltransferase 1 promotes low-grade glioma progression by activating mTORC1 signaling. Neurol Res. 2023 May;45(5):415–22. doi:10.1080/01616412.2022.2149516 PubMed PMID: 36417280.

48. Osum M, Alsaloumi L, Kalkan R. In Silico Identification of DNMT Inhibitors for the Treatment of Glioblastoma. Int J Transl Med. 2025 Dec;5(4):48. doi:10.3390/ijtm5040048

49. Pavlova NN, Zhu J, Thompson CB. THE HALLMARKS OF CANCER METABOLISM: STILL EMERGING. Cell Metab. 2022 Mar 1;34(3):355–77. doi:10.1016/j.cmet.2022.01.007 PubMed PMID: 35123658; PubMed Central PMCID: PMC8891094.

50. Li Q, Yang F, Shi X, Bian S, Shen F, Wu Y, et al. MTHFD2 promotes ovarian cancer growth and metastasis via activation of the STAT3 signaling pathway. FEBS Open Bio. 2021;11(10):2845–57. doi:10.1002/2211-5463.13249

51. Shi Y, Xu Y, Yao J, Yan C, Su H, Zhang X, et al. MTHFD2 promotes tumorigenesis and metastasis in lung adenocarcinoma by regulating AKT/GSK-3β/β-catenin signalling. J Cell Mol Med. 2021 Jul;25(14):7013–27. doi:10.1111/jcmm.16715 PubMed PMID: 34121323; PubMed Central PMCID: PMC8278097.

52. Huang M, Xue J, Chen Z, Zhou X, Chen M, Sun J, et al. MTHFD2 suppresses glioblastoma progression via the inhibition of ERK1/2 phosphorylation. Biochem Cell Biol Biochim Biol Cell. 2023 Feb 1;101(1):112–24. doi:10.1139/bcb-2022-0291 PubMed PMID: 36493392.

53. Liu X, Liu B, Wang J, Liu H, Wu J, Qi Y, et al. PHGDH activation fuels glioblastoma progression and radioresistance via serine synthesis pathway. J Exp Clin Cancer Res. 2025 Mar 19;44(1):99. doi:10.1186/s13046-025-03361-3

54. Kim D, Fiske BP, Birsoy K, Freinkman E, Kami K, Possemato R, et al. SHMT2 drives glioma cell survival in the tumor microenvironment but imposes a dependence on glycine clearance. Nature. 2015 Apr 16;520(7547):363–7. doi:10.1038/nature14363 PubMed PMID: 25855294; PubMed Central PMCID: PMC4533874.

55. Du K, Grocott L, Anichini G, O’Neill K, Syed N. Amino Acid Deprivation in Glioblastoma: The Role in Survival and the Tumour Microenvironment—A Narrative Review. Biomedicines. 2024 Oct 29;12(11):2481. doi:10.3390/biomedicines12112481 PubMed PMID: 39595047; PubMed Central PMCID: PMC11592029.

56. Tombari C, Zannini A, Bertolio R, Pedretti S, Audano M, Triboli L, et al. Mutant p53 sustains serine-glycine synthesis and essential amino acids intake promoting breast cancer growth. Nat Commun. 2023 Oct 25;14(1):6777. doi:10.1038/s41467-023-42458-1

57. Hariharan S, Whitfield BT, Pirozzi CJ, Waitkus MS, Brown MC, Bowie ML, et al. Interplay between ATRX and IDH1 mutations governs innate immune responses in diffuse gliomas. Nat Commun. 2024 Jan 25;15(1):730. doi:10.1038/s41467-024-44932-w

58. Kikuchi M, Takami H, Kobayashi Y, Nagaoka K, Kitagawa Y, Nomura M, et al. Dissecting the Immunological Microenvironment of Glioma Based on IDH Status: Implications for Immunotherapy. Cells. 2025 Jan;14(13):1035. doi:10.3390/cells14131035

59. Caverzán MD, Beaugé L, Oliveda PM, Cesca González B, Bühler EM, Ibarra LE. Exploring Monocytes-Macrophages in Immune Microenvironment of Glioblastoma for the Design of Novel Therapeutic Strategies. Brain Sci. 2023 Mar 24;13(4):542. doi:10.3390/brainsci13040542 PubMed PMID: 37190507; PubMed Central PMCID: PMC10136702.

60. Zhao W, Zhang Z, Xie M, Ding F, Zheng X, Sun S, et al. Exploring tumor-associated macrophages in glioblastoma: from diversity to therapy. NPJ Precis Oncol. 2025 May 2;9:126. doi:10.1038/s41698-025-00920-x PubMed PMID: 40316746; PubMed Central PMCID: PMC12048723.

61. Bunse L, Pusch S, Bunse T, Sahm F, Sanghvi K, Friedrich M, et al. Suppression of antitumor T cell immunity by the oncometabolite (R)-2-hydroxyglutarate. Nat Med. 2018 Aug;24(8):1192–203. doi:10.1038/s41591-018-0095-6

