## Additional File 1 for "One-Carbon Metabolic Reprogramming Stratifies Prognosis and Defines Distinct Biological States in IDH1-Mutant Gliomas"

**Research Article**

### Additional File 1

### Supplementary Figures

### Figures

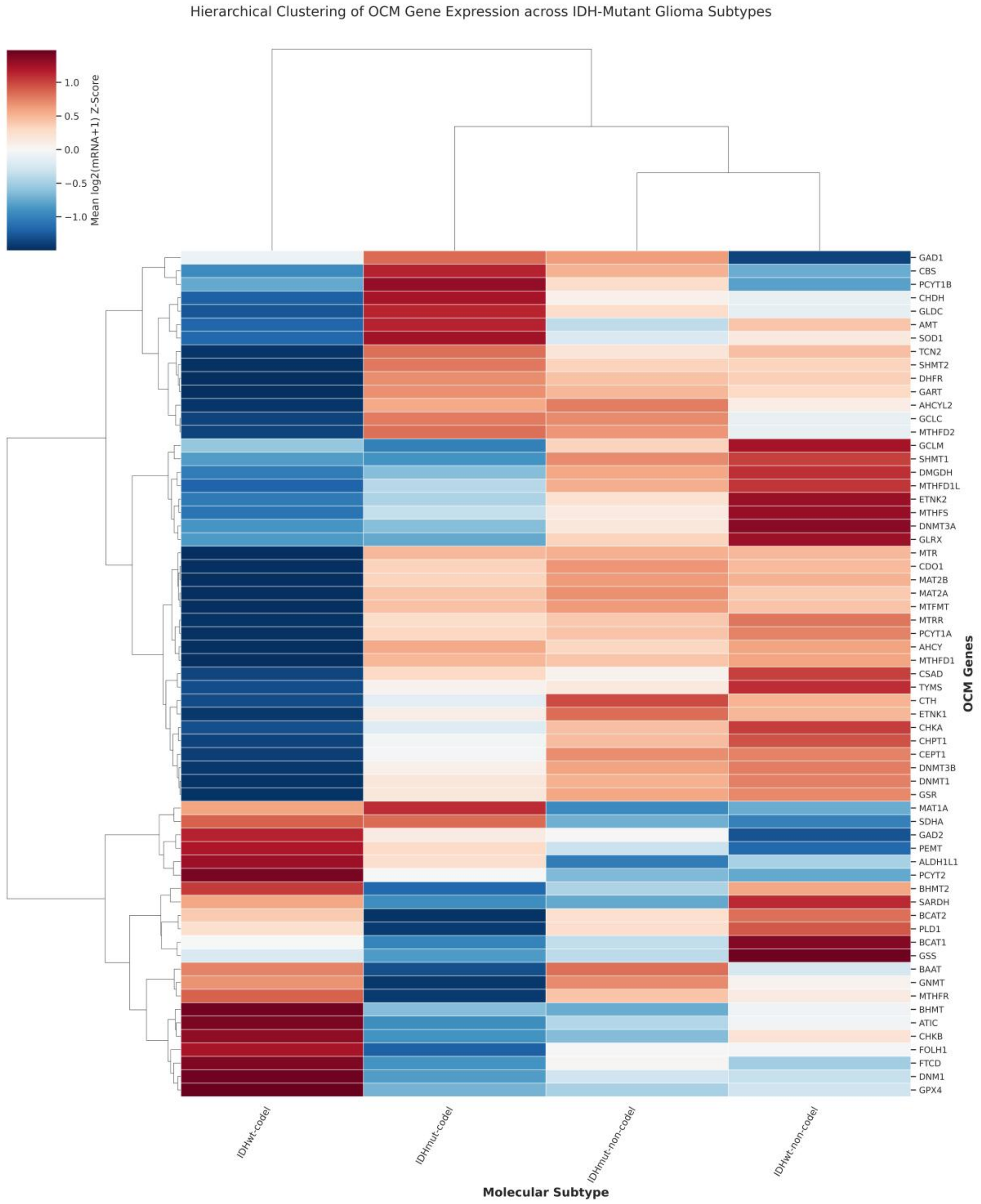


**Supplementary Figure S1: Heatmap of One-Carbon Metabolism (OCM) Gene Expression in the CGGA Cohort.** Hierarchical clustering of 64 curated OCM genes from the CGGA dataset reveals distinct transcriptional remodelling patterns segregated by IDH1 mutation status and molecular subtypes. The expression profiling visually groups the IDH-wildtype and IDH-mutant (with and without 1p/19q co-deletion) glioma phenotypes, highlighting differential metabolic dependencies and widespread suppression of proliferative OCM pathways driven by the mutant IDH1 enzyme.

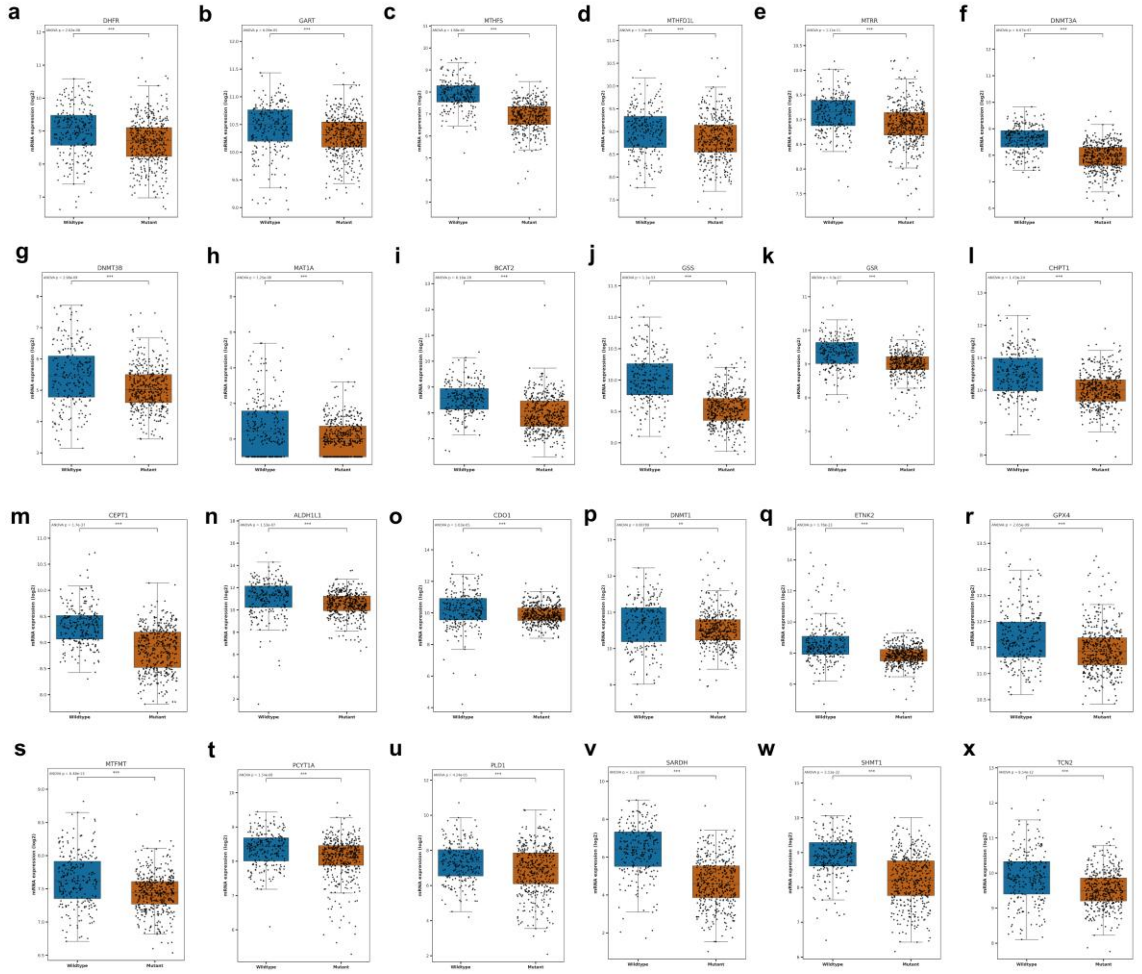


**Supplementary Figure S2: Downregulated Proliferative and Methylation OCM Markers in IDH1-Mutant Tumours (TCGA).** (a–x) Boxplots validating the significant transcriptional suppression of OCM genes associated with rapid cell growth and epigenetic maintenance in the TCGA IDH1-MT cohort compared to IDH1-WT. Key enzymes involved in *de novo* nucleotide synthesis, the DNA methylation machinery, and redox regulation exhibit markedly lower expression in IDH1-MT tumours, consistent with the slower-proliferating and less aggressive nature of this subtype.

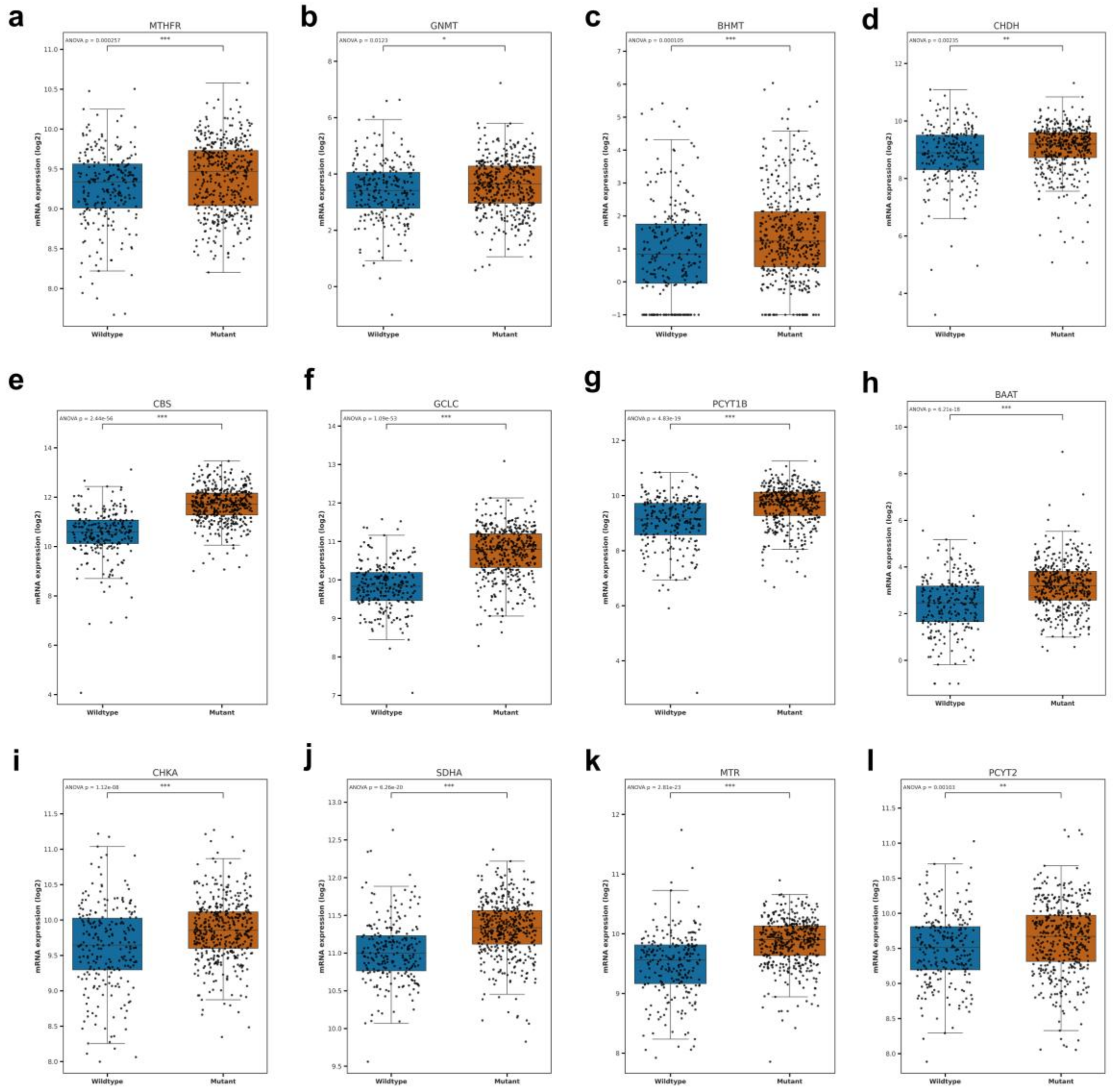


**Supplementary Figure S3: Compensatory Upregulation of Mitochondrial One-Carbon Source Pathways in IDH1-Mutant Gliomas (TCGA).** (a–l) Gene expression boxplots demonstrating significant upregulation of 1C-generating and regulatory enzymes in the TCGA IDH1-MT cohort. Enzymes driving the mitochondrial serine/glycine catabolism axis and transsulfuration pathways exhibit elevated expression, indicating a compensatory metabolic adaptation designed to sustain essential 1C flux and manage metabolic stress within the metabolically constrained IDH1-mutant environment.

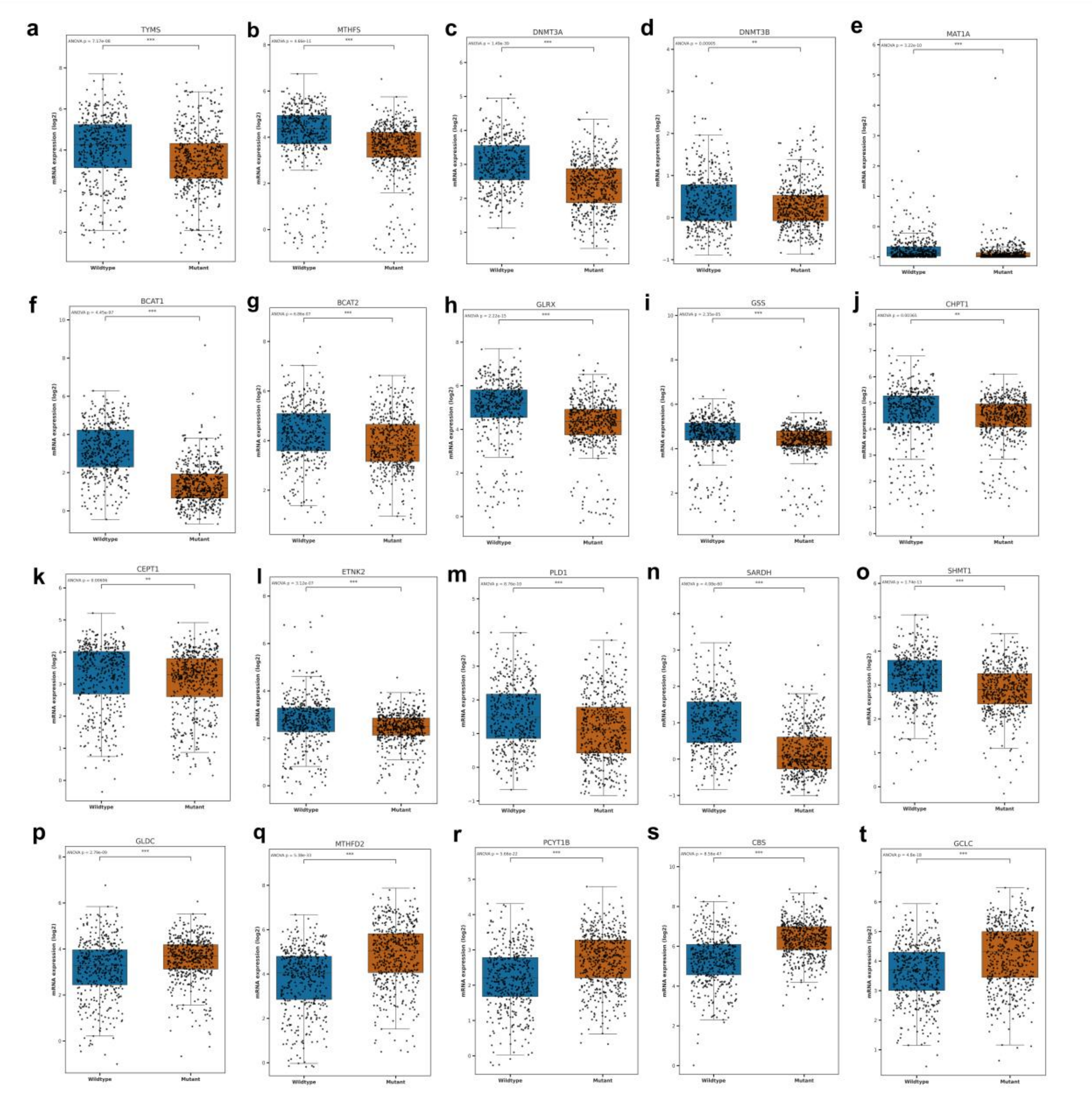


**Supplementary Figure S4: Validation of Divergent OCM Transcriptional Signatures in the Independent CGGA Cohort.** (a–t) Boxplots comparing the expression levels of key OCM genes between IDH1-WT and IDH1-MT gliomas in the CGGA dataset. The expression patterns robustly mirror the TCGA findings, confirming the broad suppression of highly proliferative markers alongside the compensatory upregulation of 1C source and maintenance genes in the IDH1-mutant phenotype.

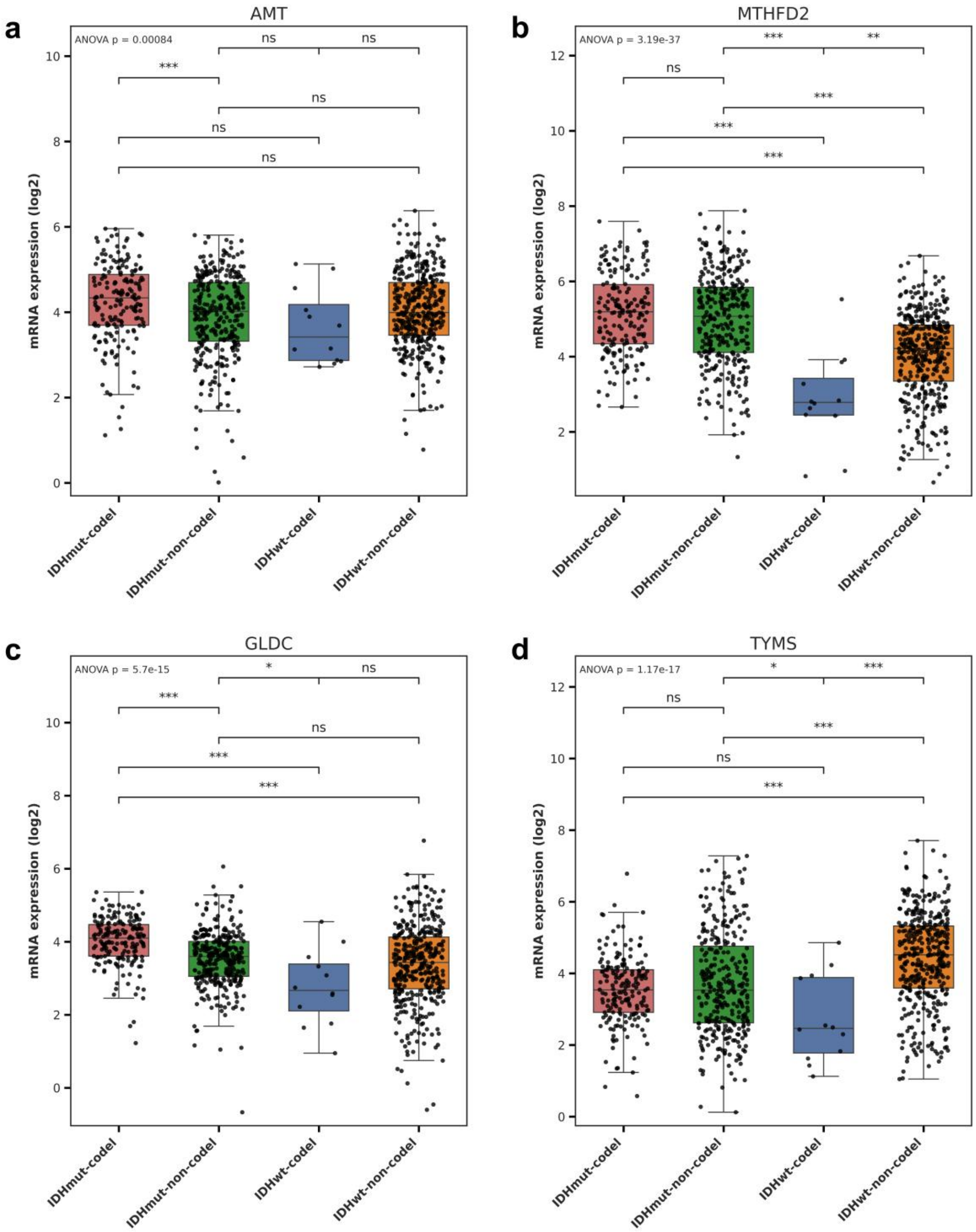


**Supplementary Figure S5: Modulatory Influence of 1p/19q Co-deletion on OCM Gene Expression.** Boxplots illustrating the specific expression profiles of key metabolic genes, a) AMT, b) MTHFD2, c) GLDC, d) TYMS, stratified across molecular subtypes, explicitly incorporating 1p/19q co-deletion status. The data indicate that co-deletion status further refines metabolic gene expression beyond that explained by IDH1 mutation status alone. In contrast, highly proliferative markers such as TYMS remain fundamentally suppressed in IDH-mutant tumours regardless of co-deletion status.

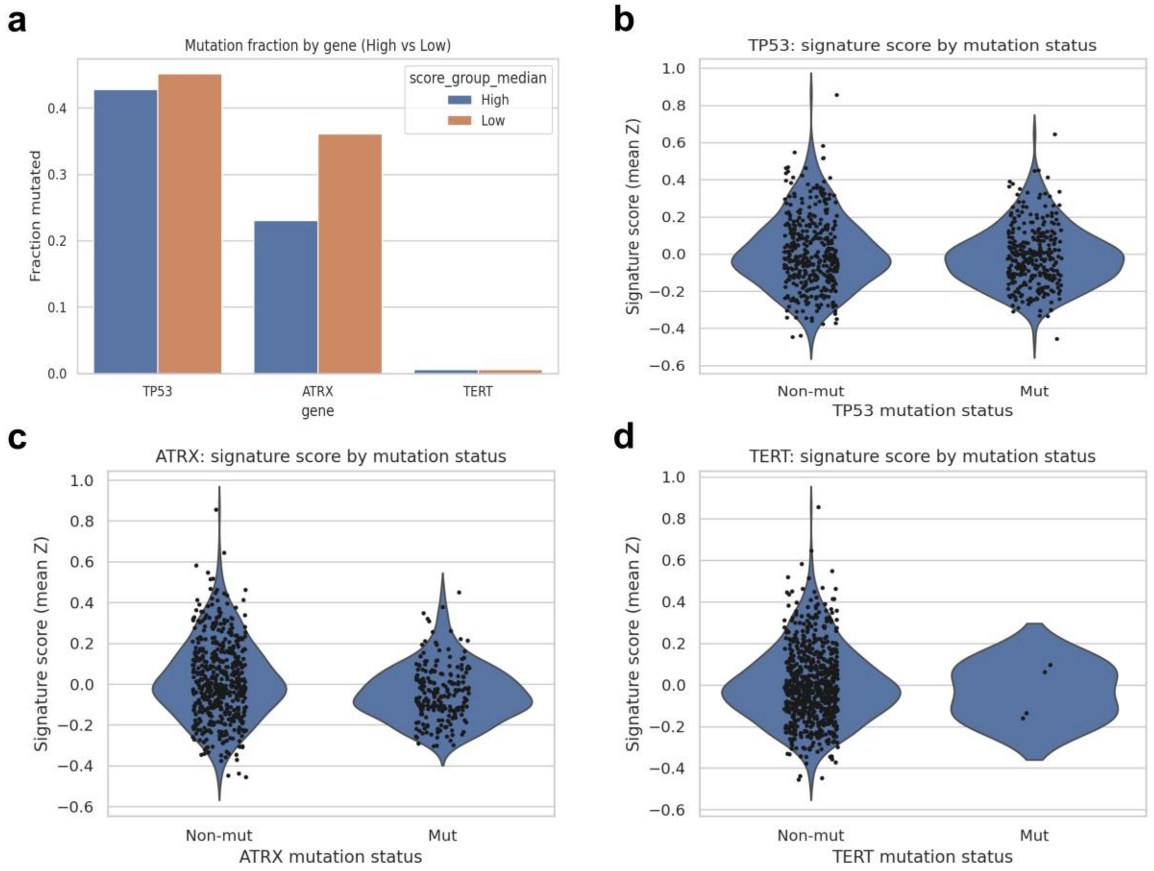


**Supplementary Figure S6: Association of OCM Signature Scores with Secondary Driver Mutations.** Analysis of the influence of TP53, ATRX, and TERT mutations on the global OCM signature. (a) Mutation fractions stratified by high and low OCM signature scores, showing TP53 as a common baseline event across groups and an enrichment of ATRX alterations specifically in the low-score group. (b-d) Continuous distribution of OCM signature scores by mutation status for TP53, ATRX, and TERT, respectively, highlighting the downward shift in overall score distribution associated with ATRX mutations, whereas TERT mutations show no clear difference and remain rare.

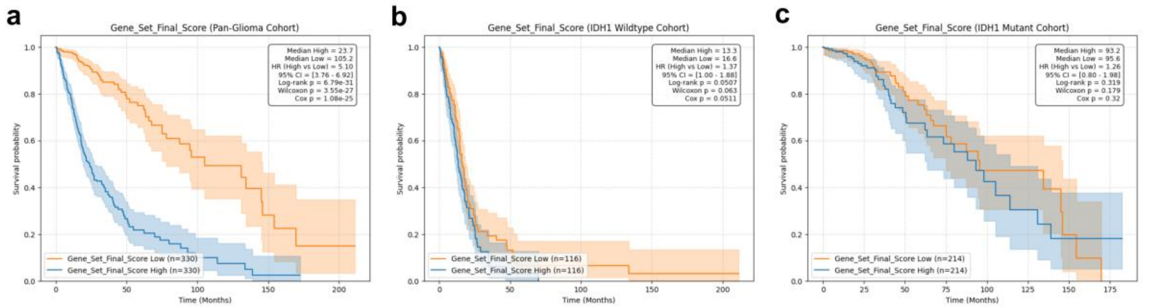


**Supplementary Figure S7: Prognostic Value of the Combined OCM Signature Score in the TCGA Cohort.** Kaplan-Meier survival analysis confirming the clinical relevance of the aggregate OCM signature. High combined OCM expression serves as a potent predictor of reduced overall survival across the Pan-Glioma cohort (a), with prognostic segregations further delineated within the highly aggressive IDH1-wildtype (b) and IDH1-mutant (c) molecular subsets.

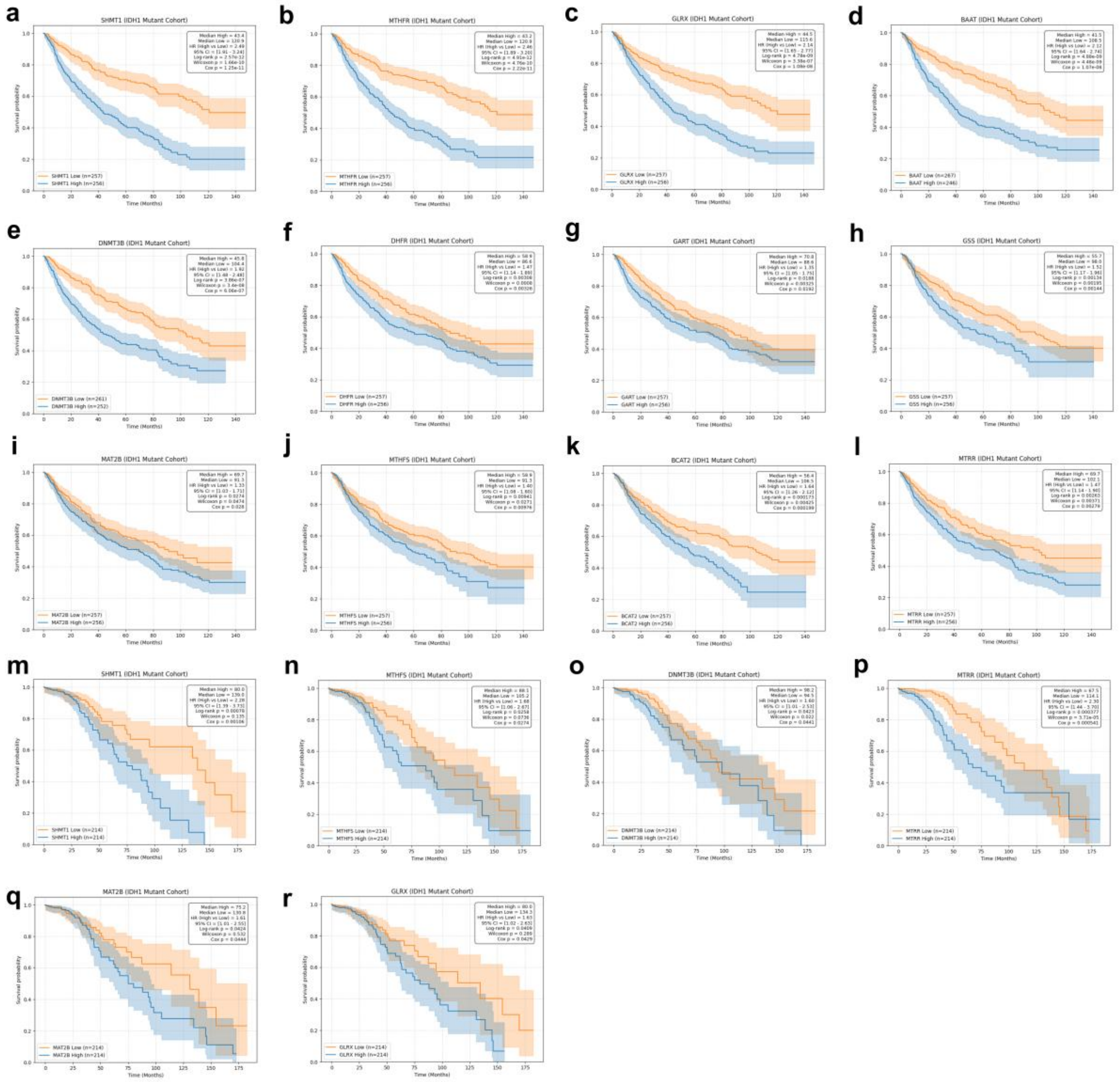


**Supplementary Figure S8: Hyperactivation of Proliferative OCM Genes Drives Poor Prognosis in IDH1-Mutant Gliomas.** Kaplan-Meier survival curves identifying detrimental OCM drivers within the IDH1-MT in CGGA (a-l), and TCGA (m-r) cohorts. High expression of specific transcriptionally suppressed genes aberrantly upregulates proliferative and methylation pathways, overriding the favourable IDH mutation status and acting as a major risk factor significantly correlated with reduced overall survival.

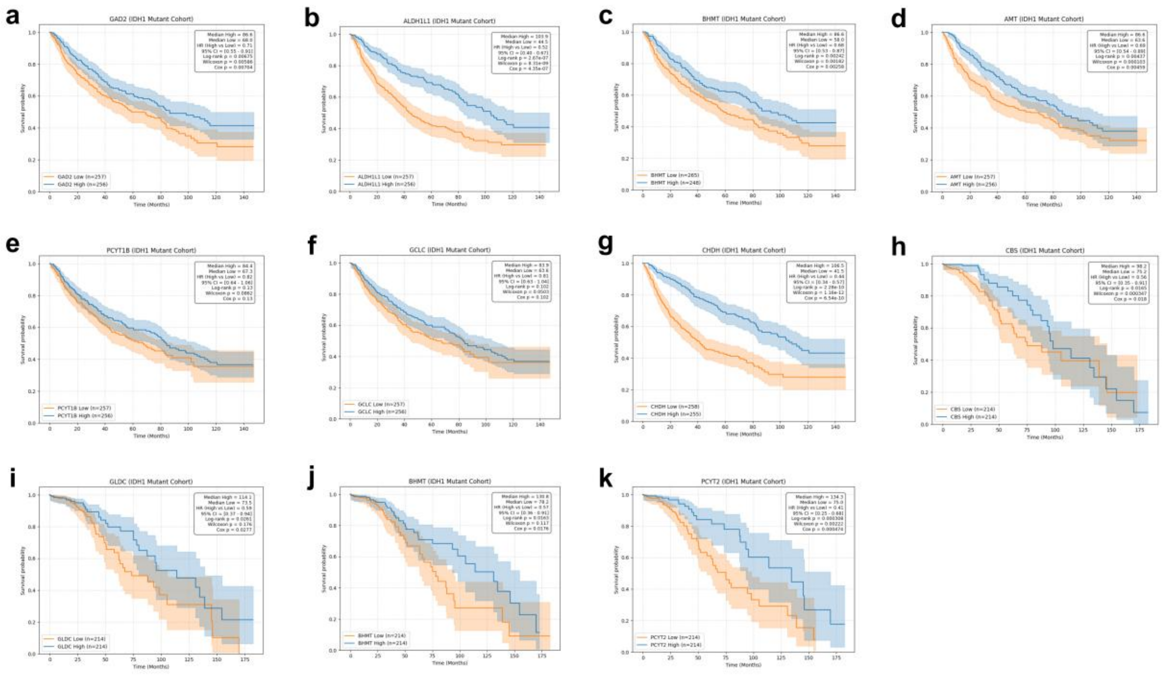

**Supplementary Figure S9: Compensatory OCM Pathway Activation Confers a Protective Survival Advantage in IDH1-Mutant Gliomas.** Kaplan-Meier plots showing protective prognostic genes within the IDH1-MT in the CGGA (a-g) and TCGA (h-k) cohorts. Elevated expression of enzymes fundamentally involved in managing 1C unit generation and lipid-membrane homeostasis confers a significant survival benefit. This illustrates that the robust efficiency of compensatory pathways is vital for maintaining the indolent tumour phenotype.

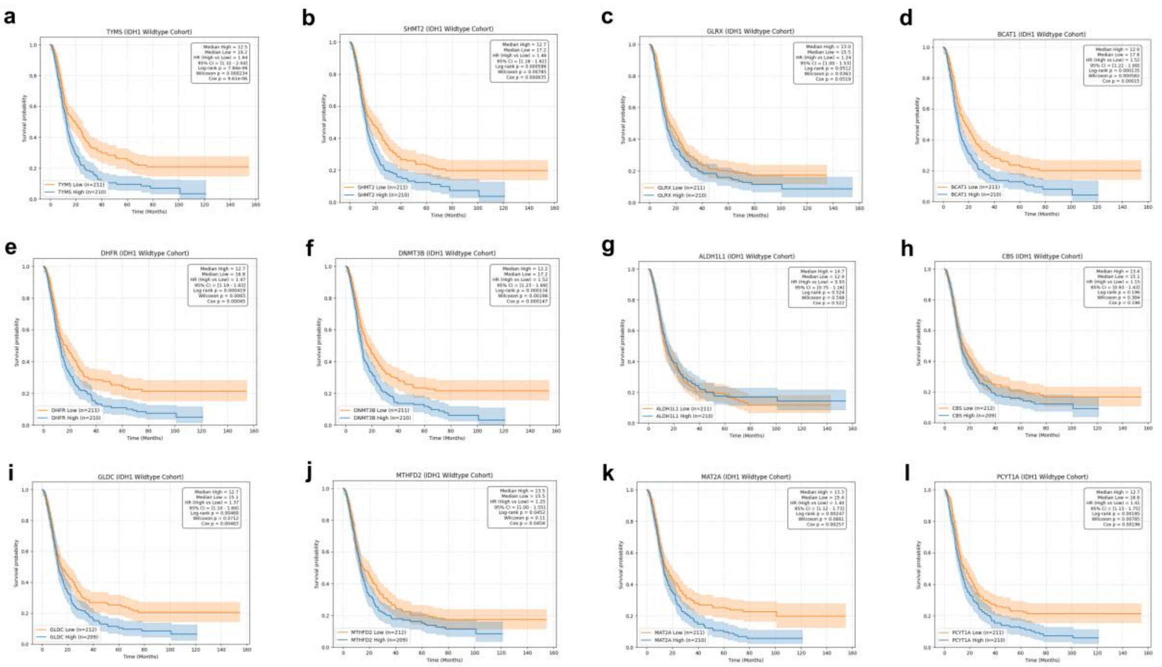

**Supplementary Figure S10: Prognostic Impact of Individual OCM Genes in IDH1-Wildtype (CGGA) Gliomas.** (a–l) Kaplan-Meier survival estimates detailing the universally detrimental effect of elevated OCM activity in the maximally aggressive IDH1-WT cohort. Unlike the divergent outcomes in IDH1-mutant disease, high expression of most OCM genes is associated with poorer survival, reflecting the sustained, high metabolic demands and inherent deleterious nature of OCM activation in this background.

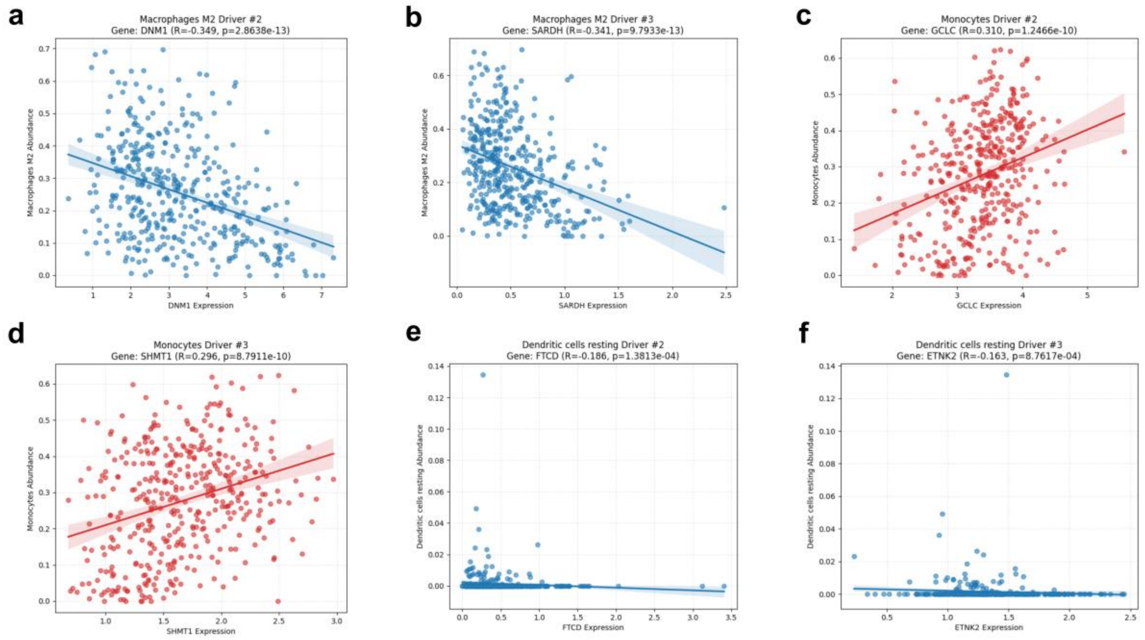


**Supplementary Figure S11: Specific OCM Genes Modulate Immune Microenvironment Interactions in IDH1-Mutant Gliomas.** Scatter plots demonstrating the significant gene-level drivers of immune cell infiltration in the IDH1-MT subset. Specific metabolic regulators correlate distinctly with immune remodelling, such as regulatory links via a) DNM1, b) SARDH, c) GCLC, d) SHMT1, as well as the negative association of e) FTCD, and f) ETNK2 with resting dendritic cells. This highlights unique, cell-specific metabolic-immune axes within the mutant tumour microenvironment.

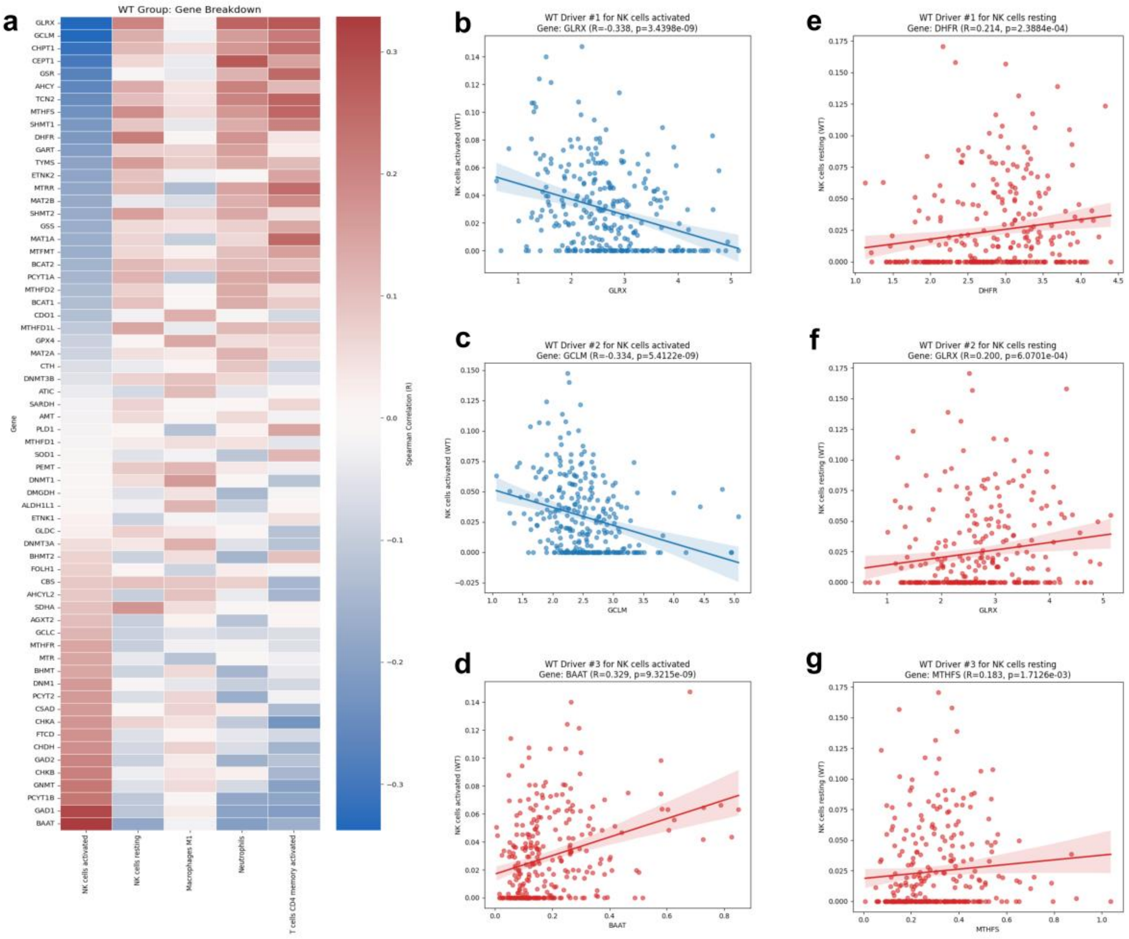


**Supplementary Figure S12: Metabolic-Immune Associations and Immune Evasion in IDH1-Wildtype Gliomas.** (a) Heatmap and (b-g) scatter plots detailing the distinct immunomodulatory profile driven by OCM genes in the IDH1-WT cohort. The data illustrate that high metabolic activity, particularly driven by redox regulators such as GLRX and GCLM, inversely correlates with activated cytotoxic NK cell infiltration, suggesting a metabolic mechanism of immune evasion specific to the wildtype landscape, in which tumour antioxidant capacity neutralises NK-mediated cytotoxicity.
