## Additional File 2 for "One-Carbon Metabolic Reprogramming Stratifies Prognosis and Defines Distinct Biological States in IDH1-Mutant Gliomas"

**Research Article**

### Additional File 2

**Supplementary Tables**

**Tables**

**Supplementary Table S1**: **The 64 One-Carbon Metabolism (OCM) Gene Signature.** A comprehensive list of the 64 core OCM candidate genes evaluated in this study, detailing their official Gene Names and corresponding Ensembl IDs. These candidate genes were strategically extracted and curated from the WikiPathways Database (pathway identifier WP2525), providing a comprehensive overview of the enzymes and transporters involved in the folate cycle, methionine cycle, and transsulfuration pathway.

| S.No. | Ensembl_ID | Gene_Name | S.No. | Ensembl_ID | Gene_Name |
|---|---|---|---|---|---|
| **1.** | ENSG00000113492 | AGXT2 | **33.** | ENSG00000159131 | GART |
| **2.** | ENSG00000101444 | AHCY | **34.** | ENSG00000001084 | GCLC |
| **3.** | ENSG00000158467 | AHCYL2 | **35.** | ENSG00000023909 | GCLM |
| **4.** | ENSG00000144908 | ALDH1L1 | **36.** | ENSG00000178445 | GLDC |
| **5.** | ENSG00000145020 | AMT | **37.** | ENSG00000173221 | GLRX |
| **6.** | ENSG00000138363 | ATIC | **38.** | ENSG00000124713 | GNMT |
| **7.** | ENSG00000136881 | BAAT | **39.** | ENSG00000167468 | GPX4 |
| **8.** | ENSG00000060982 | BCAT1 | **40.** | ENSG00000104687 | GSR |
| **9.** | ENSG00000105552 | BCAT2 | **41.** | ENSG00000100983 | GSS |
| **10.** | ENSG00000145692 | BHMT | **42.** | ENSG00000151224 | MAT1A |
| **11.** | ENSG00000132840 | BHMT2 | **43.** | ENSG00000168906 | MAT2A |
| **12.** | ENSG00000160200 | CBS | **44.** | ENSG00000038274 | MAT2B |
| **13.** | ENSG00000129596 | CDO1 | **45.** | ENSG00000103707 | MTFMT |
| **14.** | ENSG00000134255 | CEPT1 | **46.** | ENSG00000100714 | MTHFD1 |
| **15.** | ENSG00000016391 | CHDH | **47.** | ENSG00000120254 | MTHFD1L |
| **16.** | ENSG00000110721 | CHKA | **48.** | ENSG00000065911 | MTHFD2 |
| **17.** | ENSG00000100288 | CHKB | **49.** | ENSG00000177000 | MTHFR |
| **18.** | ENSG00000111666 | CHPT1 | **50.** | ENSG00000136371 | MTHFS |
| **19.** | ENSG00000139631 | CSAD | **51.** | ENSG00000116984 | MTR |
| **20.** | ENSG00000116761 | CTH | **52.** | ENSG00000124275 | MTRR |
| **21.** | ENSG00000228716 | DHFR | **53.** | ENSG00000161217 | PCYT1A |
| **22.** | ENSG00000132837 | DMGDH | **54.** | ENSG00000102230 | PCYT1B |
| **23.** | ENSG00000106976 | DNM1 | **55.** | ENSG00000185813 | PCYT2 |
| **24.** | ENSG00000130816 | DNMT1 | **56.** | ENSG00000133027 | PEMT |
| **25.** | ENSG00000119772 | DNMT3A | **57.** | ENSG00000075651 | PLD1 |
| **26.** | ENSG00000088305 | DNMT3B | **58.** | ENSG00000123453 | SARDH |
| **27.** | ENSG00000139163 | ETNK1 | **59.** | ENSG00000073578 | SDHA |
| **28.** | ENSG00000143845 | ETNK2 | **60.** | ENSG00000176974 | SHMT1 |
| **29.** | ENSG00000086205 | FOLH1 | **61.** | ENSG00000182199 | SHMT2 |
| **30.** | ENSG00000160282 | FTCD | **62.** | ENSG00000142168 | SOD1 |
| **31.** | ENSG00000128683 | GAD1 | **63.** | ENSG00000185339 | TCN2 |
| **32.** | ENSG00000136750 | GAD2 | **64.** | ENSG00000176890 | TYMS |

**Supplementary Table S2: IDH1-Specific Driver Genes.** A curated dataset identifying the 10 specific OCM genes uniquely sensitive to IDH1 mutation status. These unique candidate biomarkers were extracted by systematically comparing genes significantly associated with the IDH1 mutation with those driven by secondary TP53 and ATRX mutations using a set-based analysis, thereby isolating metabolic effectors exclusive to IDH1 biology.

| S.No. | Gene_Name |
|---|---|
| 1. | CBS |
| 2. | CHDH |
| 3. | CHKA |
| 4. | CHKB |
| 5. | CHPT1 |
| 6. | DNMT1 |
| 7. | ETNK2 |
| 8. | MAT2B |
| 9. | MTFMT |
| 10. | PCYT1B |

**Supplementary Table S3: TP53 Mutation-Associated OCM Driver Genes.** Statistical summary of the specific OCM signature genes driving mutation-associated expression shifts in relation to TP53 co-mutations. The table provides the magnitude of the transcriptional shift (Delta), the biological effect size (Cohen's d), and the corresponding p-values for all 64 genes, identifying key metabolic enzymes (e.g., MTHFR, CTH, SARDH) whose expression is significantly synergistically upregulated or downregulated by TP53 loss or mutation.

| Gene_Name | Delta | Cohen_d | p-value |
|---|---|---|---|
| MTHFR | 0.315263 | 1.190682 | 5.93E-46 |
| CTH | 0.314274 | 0.613784 | 9.95E-19 |
| SARDH | -0.3276 | -0.65197 | 7.11E-18 |
| GNMT | 0.325457 | 0.606087 | 1.09E-14 |
| BAAT | 0.069943 | 0.413034 | 3.27E-12 |
| GCLC | 0.334212 | 0.523743 | 4.53E-12 |
| ALDH1L1 | -0.49349 | -0.50025 | 3.60E-11 |
| AGXT2 | -0.01821 | -0.25125 | 5.80E-11 |
| PCYT2 | -0.20834 | -0.4778 | 1.14E-10 |
| CEPT1 | 0.145832 | 0.3989 | 2.40E-10 |
| MTHFD2 | 0.425968 | 0.471193 | 1.12E-09 |
| SHMT1 | 0.21691 | 0.432667 | 2.77E-08 |
| BCAT1 | -0.51739 | -0.42354 | 3.86E-08 |
| TCN2 | -0.2289 | -0.38403 | 1.18E-07 |
| TYMS | -0.28611 | -0.31371 | 1.57E-07 |
| DNMT3A | 0.147265 | 0.387136 | 2.08E-07 |
| GSS | -0.13466 | -0.36533 | 1.01E-06 |
| MTHFS | -0.05392 | -0.38306 | 1.91E-06 |
| MAT1A | -0.03058 | -0.23011 | 1.97E-06 |
| SDHA | -0.06844 | -0.28254 | 4.44E-06 |
| GLDC | -0.20543 | -0.29285 | 1.41E-05 |
| AHCY | -0.17463 | -0.34 | 2.02E-05 |

| | | | |
|---|---|---|---|
| FTCD | 0.133041 | 0.373136 | 2.14E-05 |
| CDO1 | 0.137752 | 0.171318 | 4.66E-05 |
| MTHFD1L | 0.120362 | 0.283783 | 6.10E-05 |
| GAD2 | -0.3693 | -0.30398 | 0.00027 |
| CSAD | -0.17802 | -0.27884 | 0.000586 |
| BHMT2 | -0.15005 | -0.26865 | 0.000587 |
| SOD1 | -0.11544 | -0.27119 | 0.000695 |
| DNM1 | -0.39325 | -0.28431 | 0.000736 |
| PEMT | -0.10555 | -0.22538 | 0.000905 |
| BCAT2 | 0.136398 | 0.242268 | 0.001447 |
| AHCYL2 | 0.154567 | 0.221128 | 0.001653 |
| GPX4 | -0.1256 | -0.2464 | 0.002056 |
| DMGDH | -0.07776 | -0.25095 | 0.002713 |
| MTRR | -0.07711 | -0.23791 | 0.003954 |
| BHMT | -0.02635 | -0.24516 | 0.013466 |
| ETNK1 | 0.073898 | 0.204757 | 0.016266 |
| MTR | 0.072575 | 0.199993 | 0.02097 |
| PLD1 | 0.061351 | 0.127854 | 0.021879 |
| MAT2A | 0.076947 | 0.132711 | 0.041611 |
| GLRX | -0.17902 | -0.22213 | 0.07232 |
| MAT2B | -0.08247 | -0.20122 | 0.126193 |
| ATIC | 0.044725 | 0.106826 | 0.131306 |
| MTHFD1 | 0.039708 | 0.112855 | 0.154443 |
| CHDH | -0.09792 | -0.14388 | 0.206672 |
| CHPT1 | -0.10384 | -0.18179 | 0.250176 |
| DNMT1 | 0.058758 | 0.117858 | 0.312887 |
| FOLH1 | -0.08221 | -0.08454 | 0.348396 |
| DHFR | -0.0488 | -0.07498 | 0.350394 |
| PCYT1B | -0.01623 | -0.02386 | 0.360295 |
| CHKA | -0.02013 | -0.04857 | 0.369484 |
| SHMT2 | -0.00172 | -0.00256 | 0.526642 |
| ETNK2 | -0.11215 | -0.1519 | 0.545749 |
| CBS | -0.02117 | -0.03943 | 0.552523 |
| CHKB | 0.007923 | 0.060845 | 0.573465 |
| PCYT1A | 0.00332 | 0.011204 | 0.575983 |
| DNMT3B | 0.037258 | 0.121628 | 0.599134 |
| GAD1 | -0.05373 | -0.04392 | 0.644134 |
| MTFMT | -0.01929 | -0.07241 | 0.677997 |
| AMT | 0.002818 | 0.062 | 0.682192 |
| GART | 0.021007 | 0.055651 | 0.701675 |
| GCLM | -0.01395 | -0.02381 | 0.839005 |
| GSR | -0.0219 | -0.06456 | 0.852308 |

**Supplementary Table S4: ATRX Mutation-Associated OCM Driver Genes.** Statistical profile outlining the OCM signature genes that significantly contribute to expression shifts driven by ATRX mutation status. The table lists the difference in mean Z-scores (Delta), effect size (Cohen's d), and statistical significance (p-values) across the OCM network. This delineates the dual regulatory role of ATRX alongside TP53, highlighting the specific genes (e.g., MTHFR, SARDH, GCLC) that are heavily modulated by ATRX genetic alterations in diffuse gliomas.

| Gene_Name | Delta | Cohen_d | p-value |
|---|---|---|---|
| MTHFR | 0.297249 | 1.076861 | 6.90E-36 |
| SARDH | -0.3677 | -0.73431 | 3.02E-20 |
| GCLC | 0.446286 | 0.711748 | 1.06E-17 |
| GNMT | 0.39407 | 0.741999 | 3.24E-17 |
| TYMS | -0.53978 | -0.60687 | 2.56E-16 |
| BAAT | 0.072504 | 0.427335 | 3.06E-16 |
| BCAT1 | -0.79134 | -0.66232 | 1.10E-13 |
| GSS | -0.2311 | -0.64299 | 2.40E-13 |
| AHCY | -0.30486 | -0.60742 | 3.28E-13 |
| MTHFD2 | 0.490995 | 0.544982 | 2.07E-11 |
| CTH | 0.248404 | 0.474829 | 2.98E-11 |
| AGXT2 | -0.02144 | -0.29615 | 2.83E-10 |
| TCN2 | -0.29501 | -0.49865 | 1.03E-09 |
| MTHFS | -0.07249 | -0.51997 | 1.17E-09 |
| AHCYL2 | 0.293602 | 0.425344 | 1.10E-07 |
| SDHA | -0.07157 | -0.29522 | 2.34E-06 |
| ALDH1L1 | -0.39086 | -0.39058 | 6.20E-06 |
| CSAD | -0.25157 | -0.39669 | 6.38E-06 |
| MAT1A | -0.03277 | -0.24659 | 2.08E-05 |
| CDO1 | 0.163032 | 0.202895 | 2.99E-05 |
| GLRX | -0.3346 | -0.42022 | 5.27E-05 |
| MTRR | -0.10303 | -0.31903 | 6.41E-05 |
| SHMT1 | 0.156107 | 0.307416 | 0.000167 |
| FTCD | 0.120744 | 0.336896 | 0.000261 |
| DNMT3B | -0.07283 | -0.23872 | 0.000291 |
| DHFR | -0.19912 | -0.30878 | 0.0003 |
| PCYT2 | -0.12733 | -0.28654 | 0.000494 |
| GAD1 | 0.342779 | 0.282448 | 0.000515 |
| GLDC | -0.14989 | -0.21244 | 0.000691 |
| ETNK1 | 0.085287 | 0.236471 | 0.004138 |
| CEPT1 | 0.061268 | 0.164853 | 0.004427 |
| GART | -0.06199 | -0.16462 | 0.004828 |
| GCLM | -0.14547 | -0.24997 | 0.00794 |
| DNMT3A | 0.065592 | 0.169833 | 0.010062 |
| PEMT | -0.07537 | -0.16037 | 0.011048 |
| BHMT2 | -0.15382 | -0.27512 | 0.011125 |
| MTHFD1L | 0.093807 | 0.220111 | 0.012596 |
| DNM1 | -0.25195 | -0.18098 | 0.012718 |
| DMGDH | -0.07779 | -0.25075 | 0.021064 |

| | | | |
|---|---|---|---|
| BHMT | -0.0199 | -0.1844 | 0.026647 |
| GPX4 | -0.09938 | -0.19428 | 0.028483 |
| SOD1 | -0.06938 | -0.16197 | 0.029041 |
| GSR | -0.07952 | -0.23567 | 0.029233 |
| MTR | 0.061876 | 0.170186 | 0.070258 |
| AMT | -0.00567 | -0.12486 | 0.098082 |
| CHDH | 0.076435 | 0.112173 | 0.109776 |
| MTFMT | -0.04433 | -0.16676 | 0.131237 |
| SHMT2 | -0.05867 | -0.08712 | 0.135549 |
| PCYT1B | 0.102193 | 0.15058 | 0.19257 |
| DNMT1 | -0.03275 | -0.06561 | 0.222053 |
| MAT2B | -0.06319 | -0.1538 | 0.22634 |
| CHPT1 | -0.08983 | -0.15703 | 0.274758 |
| GAD2 | -0.11693 | -0.09526 | 0.309789 |
| ATIC | 0.038146 | 0.091064 | 0.325838 |
| MTHFD1 | -0.01919 | -0.05449 | 0.402508 |
| PLD1 | -0.00128 | -0.00266 | 0.433666 |
| CHKA | 0.04245 | 0.102502 | 0.508841 |
| MAT2A | 0.028534 | 0.049119 | 0.529535 |
| PCYT1A | -0.01226 | -0.04137 | 0.572988 |
| BCAT2 | -0.01913 | -0.03374 | 0.645487 |
| ETNK2 | -0.13301 | -0.18024 | 0.655318 |
| CBS | -0.03319 | -0.06184 | 0.748603 |
| CHKB | -0.00333 | -0.02558 | 0.822682 |
| FOLH1 | -0.01394 | -0.01433 | 0.892977 |

**Supplementary Table S5: Tumour Immune Microenvironment Correlations in the IDH1-Mutant Cohort.** Spearman rank correlation analysis assessing the relationship between the combined OCM signature score and the estimated relative abundance of 22 distinct immune cell types within IDH1-mutant gliomas. The table details the Spearman correlation coefficients (R) and p-values, indicating a highly specific remodelling of the myeloid lineage, characterised by a strong negative correlation with immunosuppressive M2 macrophages and a significant positive correlation with monocytes.

| Immune Cell Type | Spearman Coefficient (R) | P-Value |
|---|---|---|
| Macrophages M2 | -0.22701 | 3.16E-06 |
| Monocytes | 0.185081 | 0.000155 |
| Dendritic cells resting | -0.16197 | 0.000955 |
| T cells, CD4 memory activated | -0.12728 | 0.009618 |
| T cells gamma delta | -0.10075 | 0.040718 |
| Macrophages M1 | -0.08935 | 0.069697 |
| Eosinophils | 0.07367 | 0.135009 |
| B cells memory | 0.071422 | 0.147362 |
| T cells CD4 naive | 0.069538 | 0.158363 |
| T cells CD8 | -0.05796 | 0.23989 |
| NK cells activated | 0.057399 | 0.244459 |

| | | |
|---|---|---|
| Plasma cells | -0.04446 | 0.367467 |
| Mast cells activated | 0.044207 | 0.370193 |
| T cells, follicular helper | -0.02525 | 0.608907 |
| Dendritic cells activated | 0.022159 | 0.653417 |
| Neutrophils | -0.01686 | 0.732665 |
| Mast cells resting | -0.01519 | 0.758265 |
| Macrophages M0 | 0.014192 | 0.773686 |
| B cells naive | -0.0136 | 0.782847 |
| T cells, CD4 memory resting | -0.0128 | 0.795303 |
| T cells regulatory (Tregs) | 0.001817 | 0.970636 |
| NK cells resting | -0.00052 | 0.991592 |

**Supplementary Table S6: Tumour Immune Microenvironment Correlations in the IDH1-Wildtype Cohort.** Spearman rank correlation data evaluating the immunomodulatory profile of the OCM signature score across 22 immune cell types within the IDH1-wildtype control group. Presenting correlation coefficients (R) and p-values, the table reveals a markedly distinct immune landscape compared to the mutant cohort, notably showing significant associations with activated and resting Natural Killer (NK) cells, but no significant correlation with M2 macrophages.

| Immune Cell Type | Spearman Coefficient (R) | P-Value |
|---|---|---|
| NK cells activated | -0.15633 | 0.007546 |
| NK cells resting | 0.138872 | 0.017775 |
| Macrophages M1 | 0.119878 | 0.040999 |
| Neutrophils | 0.115982 | 0.04808 |
| T cells, CD4 memory activated | 0.112636 | 0.05495 |
| T cells, CD4 memory resting | 0.11003 | 0.06085 |
| Macrophages M2 | 0.105856 | 0.071378 |
| T cells CD4 naive | -0.10166 | 0.08341 |
| Macrophages M0 | -0.08419 | 0.152 |
| Dendritic cells activated | -0.08186 | 0.163716 |
| B cells memory | 0.057123 | 0.331522 |
| T cells gamma delta | 0.052656 | 0.370784 |
| T cells, follicular helper | -0.0445 | 0.449561 |
| B cells naive | -0.04172 | 0.478332 |
| Mast cells resting | -0.04024 | 0.494177 |
| Dendritic cells resting | -0.03078 | 0.601048 |
| Mast cells activated | -0.02949 | 0.61639 |
| Monocytes | 0.014741 | 0.802277 |
| T cells regulatory (Tregs) | -0.01442 | 0.806528 |
| T cells CD8 | -0.01247 | 0.832233 |
| Eosinophils | -0.00214 | 0.971039 |
| Plasma cells | -0.00013 | 0.998277 |
